# Structural Basis of Broad Ebolavirus Neutralization by a Human Survivor Antibody

**DOI:** 10.1101/394502

**Authors:** Brandyn R. West, Anna Z. Wec, Crystal L. Moyer, Marnie L. Fusco, Phillipp A. Ilinykh, Kai Huang, Rebekah M. James, Andrew S. Herbert, Sean Hui, Ariel S. Wirchnianski, Eileen Goodwin, M. Javad Aman, Laura M. Walker, John M. Dye, Alexander Bukreyev, Kartik Chandran, Erica Ollmann Saphire

**Affiliations:** Department of Immunology and Microbiology, The Scripps Research Institute, La Jolla, CA 92037; Department of Microbiology and Immunology, Albert Einstein College of Medicine, Bronx, NY 10461; Current Address: Mapp Biopharmaceutical, San Diego, CA 92121; Department of Pathology, University of Texas Medical Branch, Galveston, TX 77555; Galveston National Laboratory, University of Texas Medical Branch, Galveston, TX 77555; Division of Virology, United States Army Medical Research Institute of Infectious Diseases, Ft. Detrick, MD 21702; Adimab LLC, Lebanon, NH 03766; Integrated Biotherapeutics, Rockville, MD 20850; Department of Immunology and Microbiology, University of Texas Medical Branch, Galveston, TX 77555; Skaggs Institute for Chemical Biology, The Scripps Research Institute, La Jolla, CA 92037

## Abstract

The structural features that govern broad-spectrum activity of broadly neutralizing, anti-ebolavirus antibodies (Abs) are currently unknown. Here we describe the first structure of a broadly neutralizing human Ab, ADI-15946, in complex with cleaved Ebola virus glycoprotein (EBOV GP_CL_). We find that ADI-15946 employs structural mimicry of a conserved interaction between the GP core and the glycan cap β17-β18 loop to inhibit infection. Both endosomal proteolysis of EBOV GP and binding of monoclonal Ab (mAb) FVM09 displace this loop, increase exposure of ADI-15946’s conserved epitope and potentiate neutralization. Our work also illuminated the determinants of ADI-15946’s reduced activity against Sudan virus (SUDV), and enabled rational, structure-guided engineering to enhance binding and neutralization against SUDV while retaining the parental breadth of activity.

**One Sentence Summary:** The first crystal structure of a broadly active antibody against surface glycoproteins of ebolaviruses identifies a highly conserved epitope beneath the glycan cap and highlights the molecular requirements for broad ebolavirus neutralization.

## Main Text

EBOV and related members of the family *Filoviridae* cause outbreaks of highly lethal disease in humans. ZMapp, an experimental cocktail of three mAbs targeting the EBOV GP spike, is currently the only Ab therapy available for emergency use against Ebola virus disease (*1*). However, the activity of ZMapp is limited to EBOV and does not extend protection to the related virulent ebolaviruses: Bundibugyo virus (BDBV) and SUDV. Characterization of known broadly neutralizing antibodies will be critical to the design of next-generation broadly protective Ab cocktails and vaccines.

The surface glycoprotein of all ebolaviruses is post-translationally processed to yield GP_1_ and GP_2_ subunits which associate into a trimer of GP_1,2_ heterodimers (*2*, *3*). The GP_1_ subunit mediates host cell attachment and receptor recognition, whereas GP_2_ mediates fusion of the virus and host membranes (*4*–*8*). The GP_2_ amino acid sequence, which includes the internal fusion loop (IFL), is relatively conserved among all ebolaviruses and is the target of pan-ebolavirus neutralizing antibodies (*9*–*11*). During infection, EBOV GP undergoes host-programmed disassembly mediated by endosomal cysteine proteases, which shed the steric bulk of the extensively glycosylated glycan cap and mucin-like domains (*12*, *13*). This in turn unmasks the receptor-binding site (RBS) within GP_1_ and reveals other previously inaccessible regions of the GP core (*8, 14*, *15*). Engagement of the intracellular entry receptor Niemann Pick C1 (NPC1) by the viral RBS leads to conformational rearrangements in GP_CL_ and culminates in virus-host membrane fusion (*8*, *15*–*18*).

The only pan-ebolavirus neutralizing Abs reported thus far target overlapping epitopes on the viral fusion loop (*9*–*11*, *19*). In contrast, the recently isolated human survivor mAb, ADI-15946, recognizes a distinct footprint in the base region of GP and potently neutralizes EBOV and BDBV; however, it lacks neutralizing and protective activity against SUDV (*9*, *19*). This property limits ADI-15946’s therapeutic usefulness as a next-generation pan-ebolavirus cocktail component (*9*, *19*). Accordingly, we sought to define the molecular determinants of ADI-15946 activity in the context of its cognate antigen, EBOV GP, and to expand its activity to SUDV.

We determined the crystal structure of ADI-15946’s fragment antigen binding (Fab) complexed with EBOV GP_CL_ to 4.1 Å resolution (Fig. 1B) (*9*). The structure was solved by molecular replacement using the previously published EBOV GP_CL_ structure as a search model (PDB: 5HJ3) and was refined to an R_work_/R_free_ of 26.2/28.1% (Table S1) (*15*). The ADI-15946 Fab targets the base of a single GP_1_/GP_2_ protomer within the GP trimer at an approximately 60° angle, with its constant domains directed downward towards the viral membrane (Fig 1B).

**Figure 1.**
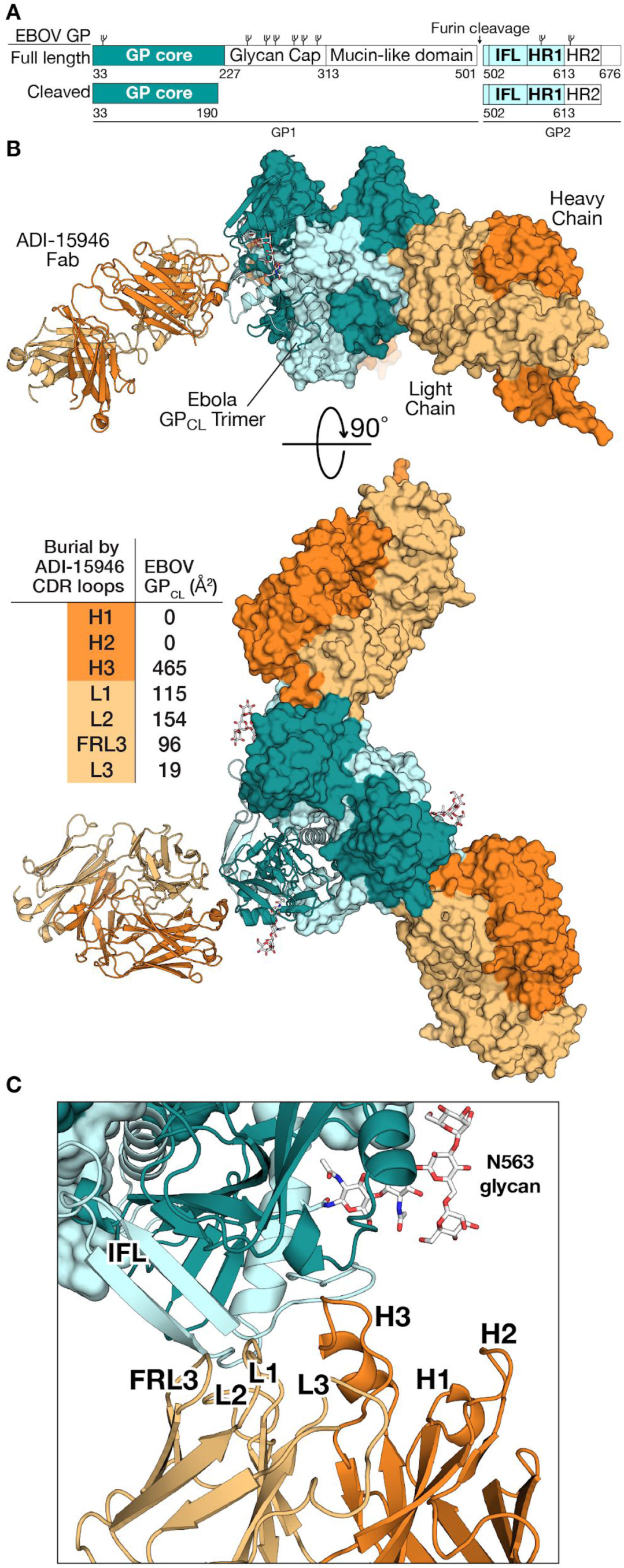
Structure of ADI-15946 in complex with Ebola virus GP_CL_. (A) The EBOV GP construct crystallized here is the ectodomain of an enzymatically cleaved form of GP (GP_CL_) resembling that generated in endosomes during viral entry lacking the glycan cap. The mucin-like domain and transmembrane regions have been deleted. (B) Crystal structure of the trimeric ADI-15946-EBOV GP_CL_ complex. GP_1_, teal; GP_2_, light blue; antibody heavy chain, orange; light chain, light orange. (C) The interaction bridges the fusion loop and other portions of GP_2_ primarily via light chain and CDR H3 contacts. CDRs H1 and H2 are not involved.

The heavy and light chains each contribute roughly half of the overall ADI-15946 binding interface: ∼55% from the heavy chain (HC) complementarity-determining region 3 (CDR H3), and the remaining 45% from the combined contributions of the light chain (LC) framework region 3 (FRL3) and CDRs 1, 2, and 3 (L1, L2, and L3 respectively) (Fig. 1B-C). Notably, CDRs 1 and 2 of the heavy chain (H1 and H2, respectively) do not participate in the Fab-GP_CL_ interface (Fig. 1B-C). Instead, HC recognition of GP_CL_ is mediated exclusively by the 22-amino-acid-long CDR H3 which buries approximately 465 Å^2^ of surface area upon binding (Fig. 1B). ADI-15946 HC contacts include the strictly conserved lysine at position 510 in the GP_2_ N terminus—where a substitution to glutamic acid, K510E, was previously shown to give rise to escape from viral neutralization (*9*). Our sidechain modeling suggests that the K510E substitution may introduce a charged/steric clash with residues D100^C^ and/or L100^F^ of CDR H3 thereby explaining the loss of antiviral activity (Fig. S1). The left boundary of ADI-15946’s footprint, formed by LC FRL3, is in immediate proximity to the β13-β14 loop that passes over the IFL and is cleaved by endosomal cathepsins during viral entry (Fig. S2) (*12*, *13*). Overlay of our structure with uncleaved GP (PDB: 5JQ3) shows that ADI-15946 binding likely impedes protease access to this loop, which is consistent with previously reported ADI-15946 inhibition of GP proteolysis (Fig. S2) (*9*).

The bottom of ADI-15946’s footprint partially overlaps those of the EBOV-monospecific binders KZ52, 2G4 and 4G7 (Fig. S3) (*2*, *20*). However, ADI-15946 likely gains cross-reactivity from an upward shift of its footprint which allows it to target a highly conserved pocket above the upper boundary of the KZ52 epitope as suggested previously (Fig. 3F, S3, and S4) (*9*). This pocket is formed by residues 71-75 of GP_1_, which fold into a 3^10^ helix, and is hereafter termed the ‘3^10^ pocket’ (Fig. 2A). The 3^10^ helix rearranges in GP_CL_ when bound to NPC1, and stabilization of this region by ADI-15946 CDR H3 may be part of the mAb’s neutralization mechanism (Fig. S4) (*8*, *9*, *15*).

**Figure 2.**
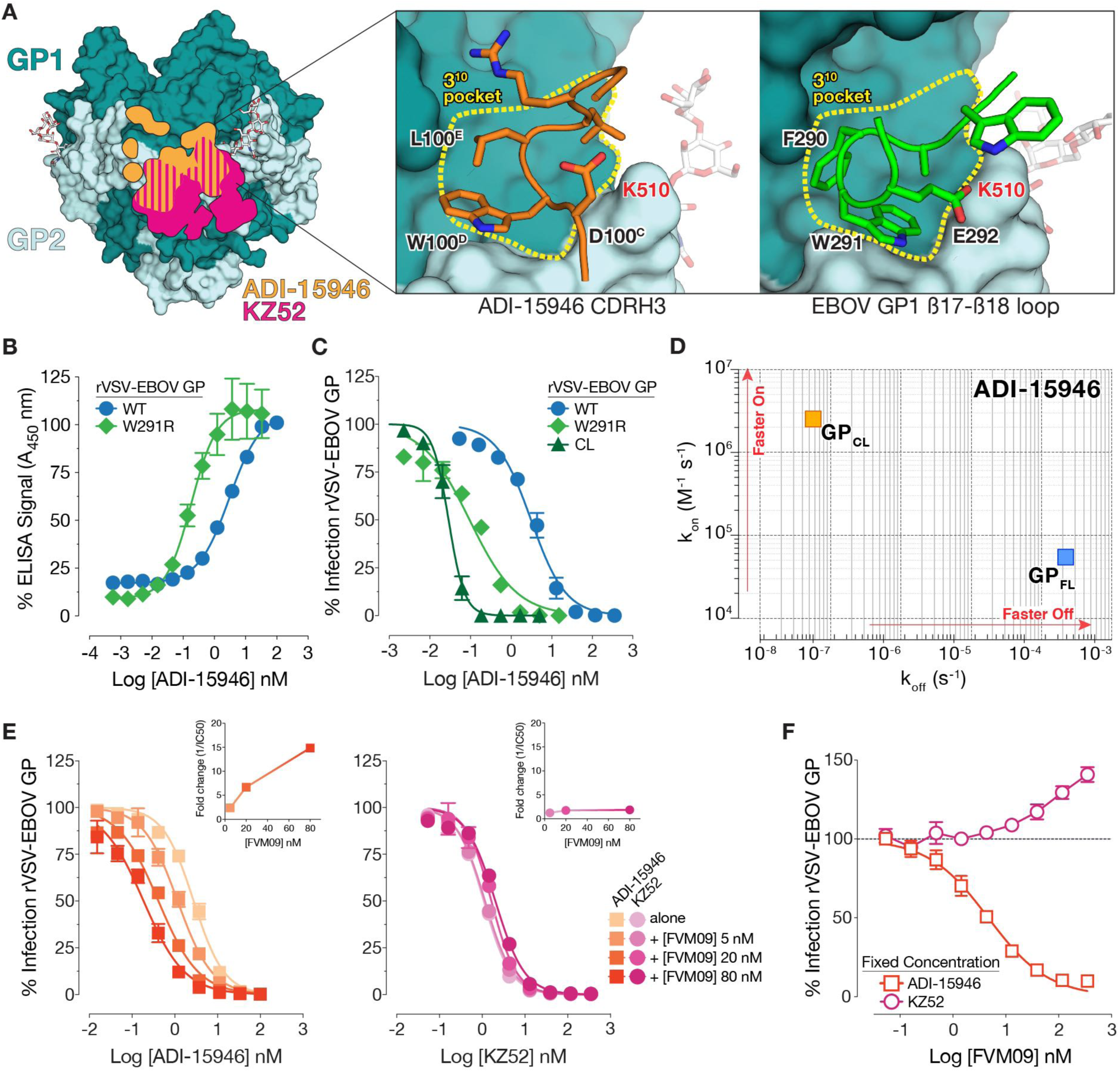
ADI-15946 binds a highly conserved epitope shielded by the mobile β17-β18 loop of the glycan cap. (A) The footprint of ADI-15946 (orange) is shifted upwards from the footprint of the EBOV-specific antibody KZ52 (pink), leaving partial overlap at the bottom of the ADI-15946 epitope (stripes). Inset, the CDR H3 of ADI-15946 (orange) binds into the 3^10^ pocket of EBOV GP (teal). This pocket is occupied in unbound GP by the β17-β18 loop (green), and in particular by residue W291, which binds into the hydrophobic pocket. (B) A point mutation in the β17-β18 loop (W291R) results in enhanced binding to EBOV GP displayed on the surface of rVSV in an ELISA. (C) The W291R point mutation in the β17-β18 loop or its proteolytic removal (CL) enhances the capacity of ADI-15946 to neutralize rVSV-EBOV GP. (D) Kinetic binding studies by biolayer interferometry reveal enhanced association rate and slower dissociation rate of ADI-15946 to GP_CL_ compared to uncleaved GP (E-F) mAb FVM09 potentiates ADI-15946 neutralization of rVSV-EBOV GP in a dose-dependent manner. (E) Addition of increasing concentrations of ADI-15946 (left), but not KZ52 (right), to fixed concentrations of FVM09 (5, 10, 20 nM) promotes rVSV-EBOV GP neutralization. The reciprocal fold change in neutralization IC_50_ is shown (inset). (F) Addition of increasing concentrations of FVM09 to a fixed, subneutralizing concentration of ADI-15946 enhances rVSV-EBOV GP neutralization. The same experiment against KZ52 actually inhibits neutralization as the concentration of FVM09 is increased.

**Figure 3.**
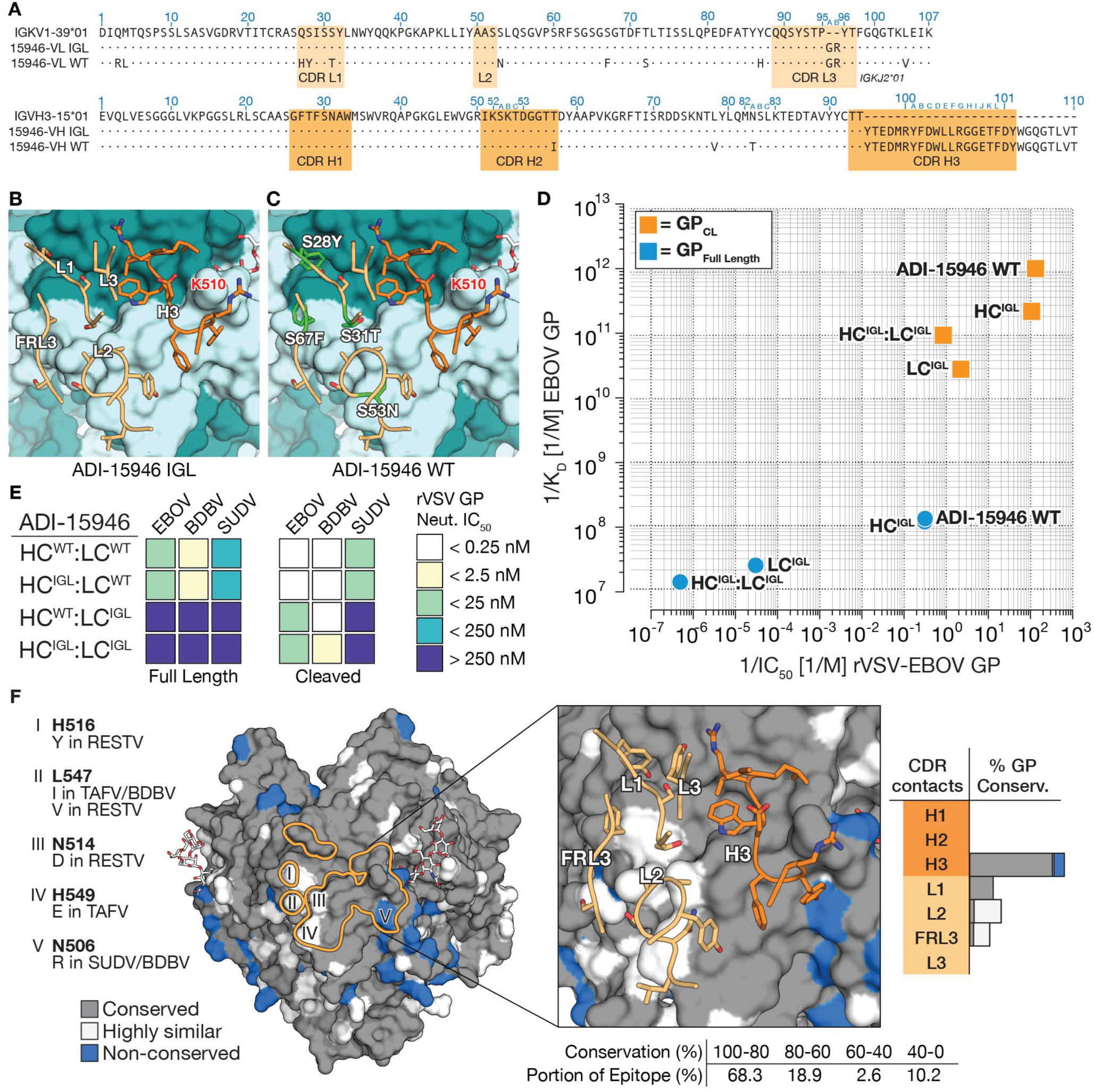
Genesis of ADI-15946. (A) Alignment of mature VH and VL sequences for ADI-15946 with their closest human germline V and J gene segments, and reconstruction of an inferred germline ancestor (IGL) bearing mature CDR-H3. Conserved residues are indicated with dots. Light and heavy chain sequences are numbered according to the Kabat scheme. (B–C) Modeling of the ADI-15946 IGL sequence (B) into the WT ADI-15946:GP_CL_ structure (C) suggests somatic hypermutations that afford enhanced recognition. (D) Comparison of the apparent equilibrium dissociation constant (1/K_D_^app^; higher value is tighter binding) for binding of ADI-15946 variants (WT, IGL, and WT:IGL chimeras) to GP_CL_ to their capacity to neutralize rVSV-EBOV GP infection (1/IC_50_; higher value is more potent neutralization). (E) Heat maps for neutralization of rVSVs bearing ebolavirus GP and GP_CL_ proteins by the indicated ADI-15946 variants. (F) Molecular surface of EBOV GP_CL_ with the ADI-15946 footprint outlined in orange. GP residues conserved across the ebolaviruses are in grey. Similar residues are in white and non-conserved residues are in blue. Differences at five sites (I-V) are listed at left. The panel at right shows the enlarged epitope with side chains of residues in ADI-15946 that contact EBOV GP_CL_ displayed as sticks.

In uncleaved GP, the 3^10^ pocket is occupied by the β17-β18 loop (GP_1_ residues 287-291) that extends down from the glycan cap (Fig. 2A and S4) (*8*, *21*). Interactions of ADI-15946 CDR H3 residues D100^C^, W100^D^, and L100^E^ with the 3^10^ pocket mimic the contacts made by β17-β18 loop residues E292, W291, and F290, respectively (Fig. 2A). Correspondingly, we postulated that the viral β17-β18 loop peptide competes for binding with CDR H3 of ADI-15946 in full length GP (GP_FL_) (Fig. 2A, S4, and S5). To test the effect of β17-β18 loop removal on ADI-15946 binding and neutralization, we measured the apparent binding constant (K_D_^app^) for ADI-15946 interaction with EBOV GP_FL_ and GP_CL_ via Bio-Layer Interferometry (BLI) and its neutralization of recombinant vesicular stomatitis virus (rVSV) particles bearing EBOV GP_FL_ and GP_CL_ (rVSV-EBOV GP_FL_ and rVSV EBOV GP_CL_, respectively). We observed a 10,000-fold improvement in binding and a 100-fold improvement in neutralization potency against EBOV GP_CL_ compared to full-length GP (Fig. 2C, 3D-E, and Tables S2 and S4). Comparison of association (k_on_) and dissociation (k_off_) rates of ADI-15946 to EBOV GP_FL_ and GP_CL_ showed that the improved binding to GP_CL_ is primarily driven by over 1000-fold slower k_off_ and a modest improvement of the association rate (10-fold increased k_on_) (Fig. 2D, S16 and table S2). Given the sizeable contribution of CDR H3 to the binding interface, the slower dissociation rate against GP_CL_ likely results from its unobstructed access to the 3^10^ pocket.

To specifically probe the importance of the β17-β18 loop’s hydrophobic packing into the 3^10^ pocket to the shielding of the ADI-15946 epitope, we tested ADI-15946 binding and neutralization against rVSVs bearing an EBOV GP variant with an arginine substitution at position 291 (rVSV-EBOV GP_W291R_). Modeling of the arginine sidechain suggested that the W291R substitution would displace the β17-β18 loop from the conserved pocket by introducing charged and steric clashes with the GP_2_ residue asparagine 512 (Fig. S6). Consistent with this hypothesis, we observed a 10-fold enhancement in ADI-15946 binding to EBOV GP_W291R_ compared to wild-type EBOV GP (GP_WT_) presented on rVSV and a >10-fold increase in neutralization efficiency of rVSV bearing EBOV GP_W291R_ relative to EBOV GP_WT_ (Fig. 2B-C). The intermediate neutralization potency of rVSV-EBOV GP_W291R_ compared to GP_CL_ and GP_WT_, suggests that displacement of the β17-β18 loop from the 3^10^ pocket is the limiting factor for binding of ADI-15946 to EBOV GP (Fig. 2C).

Previous work suggests that a non-neutralizing mAb, FVM09, recognizes and “peels away” the β17-β18 loop from the base, thereby enhancing neutralization by the base-binding mAb 2G4 (*22*, *23*). Given the more profound role of the β17-β18 loop in restricting access of ADI-15946 to its base epitope (see above), we evaluated the hypothesis that FVM09 could also work in concert with ADI-15946 to neutralize virus. We first tested competition between ADI-15946 and FVM09 using BLI-based competitive binding assays and found that both mAbs could bind to EBOV GP simultaneously (Fig. S7). We next tested the effect of FVM09 on ADI-15946 neutralization of rVSV-EBOV GP or authentic EBOV in the presence of three fixed concentrations of FVM09. FVM09 potentiated ADI-15946 neutralization in a concentration-dependent manner by >10-fold, but had no effect on neutralization by KZ52 (Fig. 2E). Conversely, titration of FVM09 into a constant, sub-neutralizing concentration of ADI-15946 also showed dose-dependent enhancement of ADI-15946 neutralizing activity (Fig. 2F). In contrast, KZ52 neutralization was somewhat reduced in the same assay, perhaps because KZ52 binds along the surface of the β17-β18 loop whereas ADI-15946 binds beneath the β17-β18 loop (Fig. 2F, S8 and S9). We observed similar neutralization enhancement trends with authentic EBOV (Fig. S9A-B), and to a lesser extent, with rVSV-SUDV GP (Fig. S9C-D). The functional significance of the strict conservation of the amino acid sequence in the 3^10^ pocket and the β17-β18 loop among ebolavirus GPs remains unknown, but it may relate to the transduction of conformational changes in GP upon NPC1 binding (*8*). Our results demonstrate that rare neutralizing antibodies like ADI-15946 can access and exploit this site for broad ebolavirus neutralization, presumably by neutralizing its functional role in viral entry. The enhancement of ADI-15946 binding and neutralization in the presence of β17-β18 loop binders such as FVM09 may reflect a mechanism by which specific combinations of glycan cap- and base-binding mAbs can synergize to interdict viral infection during a natural polyclonal immune response.

To delineate the molecular basis of ADI-15946’s broad activity, we assigned its light and heavy chain variable domain (VL and VH, respectively) sequences to their most probable inferred germline progenitors (IGL) (Fig. 3A). We found that reversion of three residues introduced by somatic hypermutation (SHM) in the VH to germline (LC^WT^:HC^IGL^; CDR H3 is retained fully mature) had no appreciable impact on binding of full-length or cleaved EBOV, BDBV, or SUDV GP or neutralization of rVSV bearing EBOV, BDBV, or SUDV GP (Fig. 3D-E, and Fig. S10). These findings agree with our structural observations, since only CDR H3 participates in binding to EBOV GP_CL_ (Fig. 1B-C). The structure also suggests that the length of CDR H3 is critical for the mAb’s ability to access the 3^10^ pocket. We tested three truncation mutants of ADI-15946 CDR H3 and found complete loss of binding and neutralization with all three (Fig. S11).

Reversion of ten SHM-introduced residues in the VL outside of CDR L3 (LC^IGL^:HC^WT^; residues 3, 4, 27, 28, 31, 53, 67, 72, 87, 104) had a profound impact on neutralization of rVSVs bearing EBOV, BDBV, or SUDV GP (Fig. 3A, and S9). Both the LC^IGL^:HC^WT^ and the combined LC^IGL^:HC^IGL^ Ab variants were non-neutralizing against rVSV-EBOV GP_FL_ despite retaining a fully mature CDR H3 and CDR L3, suggesting that LC contacts outside of CDR L3 make key contributions to GP recognition (Fig. 3E). The LC^IGL^:HC^WT^ and the LC^IGL^:HC^IGL^ germline-approximating Ab variants retained some neutralizing activity against EBOV GP_CL_ and BDBV GP_CL,_ likely due to improved access of CDR H3 to the 3^10^ pocket (Fig. 3E, and Table S2). However LC^IGL^:HC^IGL^ did not neutralize rVSV-SUDV GP_CL_, probably due to clashes between ADI-15946 CDR H3 and residues in the N-terminal tail of SUDV GP_2_ (Fig. S10). Our germline-approximating variants LC^IGL^:HC^WT^ and the LC^IGL^:HC^IGL^ neutralized the rVSV-EBOV GP_W291R_ variant more potently than rVSV-EBOV GP_CL_ which suggests that additional LC contacts that are absent in GP_CL_, likely between the Ab and the glycan cap, contribute to ADI-15946’s activity against GP_FL_ (Fig. S2, S10G and Table S2).

Our analysis of ADI-15946’s epitope conservation shows that the limited SUDV reactivity and lack of activity against Reston virus (RESTV) likely arise from the amino acid sequence divergence on the edges of its footprint (Fig 3F, S12, and S13). There are only five non-conserved residues across all five ebolavirus GPs involved in the interface with ADI-15946, all located in GP_2_: N506, N514, H516, L547, and H549 (Fig. 3F and Fig. S12). These are predominantly contacted by CDR L2 and FRL3 (Fig. S12). However, CDR H3 contacts one key residue in EBOV GP (Asn 506) that is not conserved in SUDV GP (Arg 506). Using this information, we separately mutated nearby residues in ADI-15946 CDR H3, R100 and Y100^A^, to alanine to reduce steric and charge clashes with SUDV R506. The ADI-15946 mutant Y100^A^ showed decreased binding to both SUDV and BDBV GP compared to wild-type ADI-15946 (Fig S14). However, the mAb variant called 46M1, containing R100A, showed increased binding to SUDV GP ectodomain by ELISA and slightly improved neutralization of authentic virus compared to the parental antibody (Fig. 4E-L and S13).

**Figure 4.**
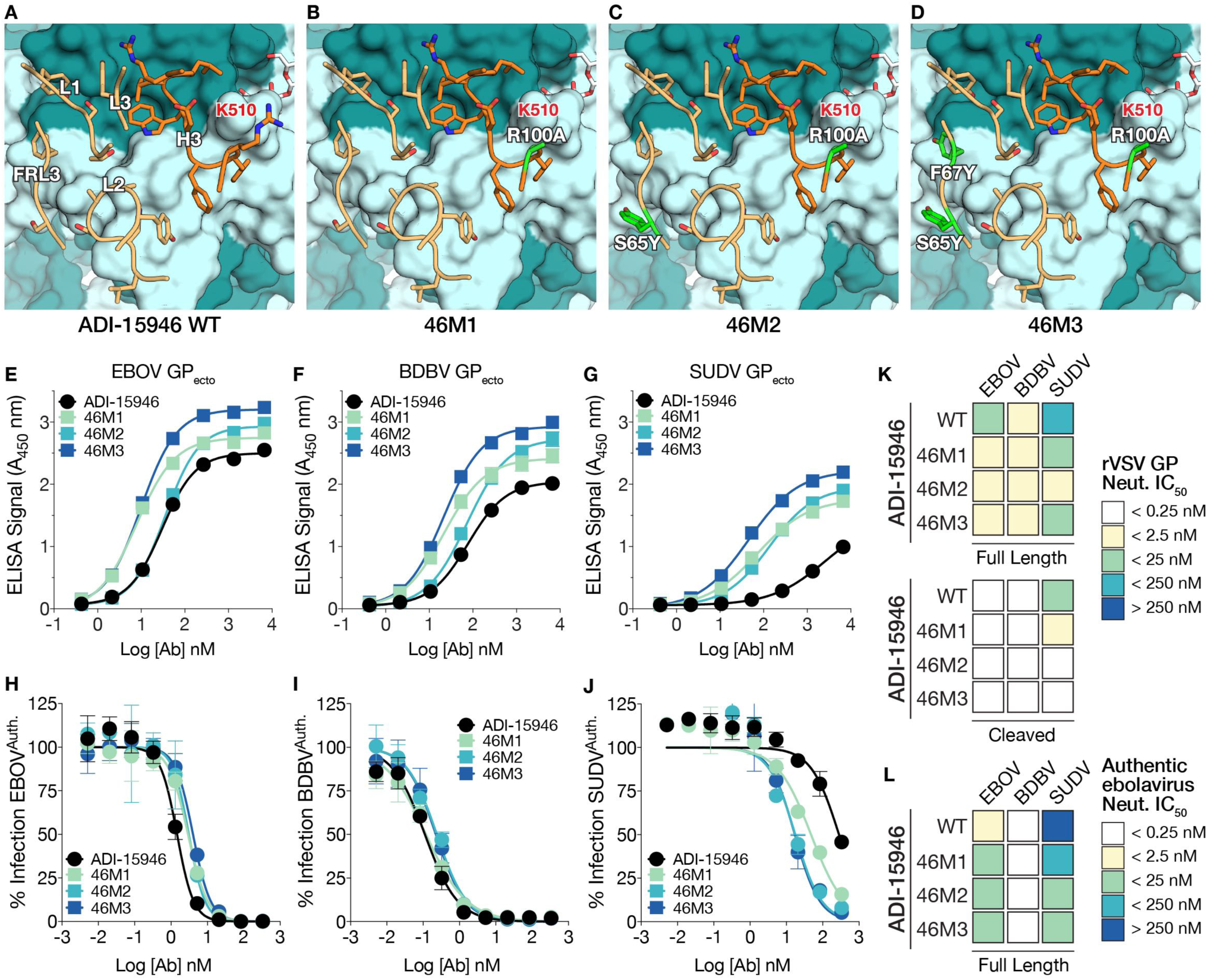
Structure-guided affinity maturation of ADI-15946. (A) Crystal structure of ADI-15946 in complex with GP_CL_. CDRs are illustrated in dark orange for the heavy chain and light orange for the light chain respectively. (B–D) Molecular models of the 46M1, 46M2, and 46M3 mutants respectively. Engineered side chains that differ from wild-type are colored in green. (E–G) Binding of recombinant EBOV, BDBV, and SUDV GP ectodomains by the indicated ADI-15946 variants determined by ELISA. (H–J) Neutralization of authentic EBOV, BDBV and SUDV by the indicated ADI-15946 variants. (K–L) Heat maps for neutralization potency (IC_50_) of each ADI-15946 variant against rVSVs (K) and authentic filoviruses (L). In panel K, neutralization of rVSVs bearing full-length GPs and cleaved GPs is shown at the top and bottom, respectively.

Structural alignment of additional Fab-GP complexes enabled comparison of the binding features of ADI-15946 with monospecific mAbs KZ52 and c4G7, which share part of their footprints with ADI-15946. (Fig. S3, S8, and S15) (*2, 20*). We observed that the HC of c4G7 places a tyrosine residue in a similar position and orientation to that of the ADI-15946 LC residue F67 and another tyrosine residue proximal to ADI-15946 LC residue S65 (Fig. S15). We then attempted to mimic this double tyrosine motif by incorporation of light chain S65Y and F67Y mutations into ADI-15946 to enhance binding to SUDV GP. The construct bearing heavy chain R100A and light chain S65Y is called variant 46M2, and ADI-15946 bearing the R100A/S65Y pair plus LC substitution F67Y is called variant 46M3. The 46M2 and 46M3 antibodies showed enhanced binding of SUDV, while maintaining parental binding to EBOV and BDBV GPs (Fig. 4E-G). 46M2 and 46M3 also exhibited a 16- to 33-fold improvement in their capacity to neutralize SUDV GP-bearing rVSV and authentic SUDV respectively, while retaining neutralization of EBOV and BDBV (Fig. 4H-L and S13). We show that relatively few mutations introduced in the 46M2 and 46M3 variants resulted in improved binding and neutralization of SUDV and confirmed the importance of the LC FR3 region to the binding interface.

No high-resolution structure of a broadly neutralizing antibody in complex with an ebolavirus glycoprotein has yet been reported. The crystal structure of ADI-15946 Fab in complex with EBOV GP_CL_ and accompanying biochemistry presented here illuminate antigenic features of GP that dictate access to a broadly conserved but previously unappreciated epitope shared by ebolaviruses. This site of vulnerability is shielded by the descending β17-β18 loop of the glycan cap, removal of which potentiates ADI-15946 activity. We also uncover potentially synergistic neutralization between ADI-15946 and the non-neutralizing glycan cap binder FVM09. Our findings support a mechanism of action whereby ADI-15946 gains enhanced neutralizing activity against the endosomal, cleaved viral GP species generated by host proteases during entry. In contrast, other potent neutralizers that target the GP base, such as KZ52 and the ZMapp components c2G4 and c4G7, have been shown to lose their activity against GP_CL_ (*9*). The mechanism of neutralization by ADI-15946 may also involve inhibition of conformational changes required for membrane fusion by anchoring to the GP_1_-GP_2_ interface across the IFL thereby preventing its unraveling from the GP_1_ core during fusion triggering. In addition, binding of ADI-15946 in the 3^10^ pocket could prevent rearrangement of GP_1_ residues 71-75 upon binding of GP_CL_ to NPC1 by mimicking the hydrophobic interactions of the β17-β18 loop that packs into the pocket in GP_FL_ (Fig. S4) (*8*). Rational structure-guided substitutions, two in the LC FRL3 and one in CDR H3, enhanced both binding and neutralization activity against SUDV without loss of activity against EBOV or BDBV.

Analysis of germline-reverted versions of the antibody indicated that a limited number of somatic mutations were required to bind this conserved GP site and that many of these mutations occurred in the framework region. Klein *et al.* show that somatic mutations in the immunoglobulin framework region enhance affinity by decreasing the dissociation rate and are generally necessary to achieve broad neutralization of HIV-1 (*25*). We found that, similar to broadly neutralizing antibodies against HIV-1, ADI-15946 also required somatic maturation of the framework region, which decreased its dissociation rate, in order to achieve broad neutralization of ebolaviruses (*25*).

Our work details structural features of GP recognized by a broadly active anti-ebolavirus antibody and provides specific strategies (i.e. cleaving GP to remove the glycan cap or recombinantly expressing a Δβ17-β18 loop variant of GP) to enable the targeting of a highly conserved site in the design of vaccines and immunotherapeutics. The structure also provided the roadmap for rational engineering of the parent Ab to enhance its activity against Sudan virus. The resulting broadly neutralizing antibody may be suitable as a component of a protective cocktail targeting the three ebolaviruses known to cause human disease outbreaks.

## Acknowledgements

We acknowledge National Institutes of Health grants U19 AI 109762 (EOS, KC, JMD), U19 AI109711 (AB), Defense Threat Reduction Agency HDTRA1-13-1-0034 (AB), R01 AI126587(MJA), and the Viral Hemorrhagic Fever Immunotherapeutic Consortium for support. Use of the Stanford Synchrotron Radiation Lightsource, SLAC National Accelerator Laboratory, is supported by the U.S. Department of Energy, Office of Science, Office of Basic Energy Sciences under Contract No. DE-AC02-76SF00515. The SSRL Structural Molecular Biology Program is supported by the DOE Office of Biological and Environmental Research, and by the National Institutes of Health, National Institute of General Medical Sciences (including P41GM103393). Opinions, conclusions, interpretations, and recommendations are those of the authors and are not necessarily endorsed by the U.S. Army. The mention of trade names or commercial products does not constitute endorsement or recommendation for use by the Department of the Army or the Department of Defense. This is manuscript number 29630 of The Scripps Research Institute. Coordinates and structure factors have been deposited in the Protein Data Bank (PDB:XXXX).

## Supplementary Materials

### Materials and Methods

#### Cloning

Mutants were introduced into parental 15946 by site-directed mutagenesis using Quik-Change (Agilent), with all mutations confirmed by sequencing (Eton).

#### Protein Expression & Purification

Expression and purification of EBOV GP_CL_ was performed as described previously (*15*). Briefly, Ebola virus GP (lacking the mucin domain residues 312 to 462) was produced by stable expression in Drosophila melanogaster S2 cells. Effectene (Qiagen) was used to transfect S2 cells with a modified pMT-puro vector plasmid containing the GP gene of interest, followed by stable selection of transfected cells with 6 μg/ml puromycin. Cells were cultured at 27°C in complete Schneider’s medium for selection and then adapted to Insect Xpress medium (Lonza) for large-scale expression in 2-liter Erlenmeyer flasks. Secreted GP ectodomain expression was induced with 0.5 mM CuSO4, and supernatant harvested after 4 days. Ebola virus GP was engineered with a double Strep-tag at the C terminus to facilitate purification using Strep-Tactin resin (2-1201-010) (Qiagen) and then further purified by Superdex 200 (GE) size exclusion chromatography (SEC) in 10 mM Tris-buffered saline (Tris-HCl, pH 7.5, 150 mM NaCl [TBS]). EBOV GP_CL_ was produced by incubation of 1 mg GP with 0.02 mg thermolysin overnight at room temperature in TBS containing 1 mM CaCl2 and purified using Superdex 200 SEC.

ADI-15946 Fab used for crystallization experiments was cloned into a modified pMT-puro vector with a heavy chain C-terminal Strep-tag, and then expressed and purified according to the protocol for GP_CL_ with the exception that SEC was performed with a Superdex 75 column (GE) (*15*).

ADI-15946 IgG used for ELISA and neutralization assays were produced in ExpiCHO cells (ThermoFisher Scientific) and purified via Protein A chromatography according to the standard ThermoFisher ExpiCHO protocol for a 25 mL culture volume. Cells were pelleted 8 days post transfection by centrifugation for 30 minutes at 3000 xg and supernatant was collected for purification of soluble ADI-15946 IgG. Supernatant was flowed over 2 mL of Pierce Protein A Plus Agarose resin (ThermoFisher Scientific). Column was washed with 5 column volumes of DPBS (Gibco) and then IgG was eluted with DPBS supplemented with 25 mM glycine pH 2.2.

#### Crystallography & Structure Determination

Trimeric EBOV GP_CL_ was complexed with ADI-15946 Fab fragments, and the resulting complex was then purified via SEC. The purified EBOV GP_CL_-ADI-15946 Fab complex was concentrated to 4.2 mg/ml in TBS. The crystal drops consisted of a 1:1 ratio of protein/well solution. Crystals grew over the course of one month in 0.2 M sodium citrate tribasic dihydrate pH 8.2 and 20% polyethylene glycol 3350. Crystals were cryoprotected with 20% glycerol and flash frozen in liquid nitrogen for storage and shipping. Diffraction data was collected remotely on SSRL beamline 12-2 on a pilatus 6M detector (*26*–*29*). Data was processed using XDS (*30*, *31*), and the structure was determined using molecular replacement with PHASER (*32*), within the CCP4 suite (*33*), using the structure of EBOV GP_CL_ (PDB: 5HJ3) as an initial search model (*15*). Iterative rounds of model building were performed using Coot (*34*), and each round was refined with Phenix (*35*). Five percent of the data was set aside prior to refinement for the R_free_ calculations for each data set (*36*). The statistics and stereochemistry of the crystal structure were checked using the MolProbity server (*36*, *37*). Structural figures were rendered using Open Source PyMOL (PyMOL Molecular Graphics System, version 1.7.0.0; Schrödinger, LLC).

Structural Alignment and Visualization of Ebolavirus Glycoproteins

Alignment was performed using clustalomega on uniprot with the following protein sequences: Zaire ebolavirus: Q05320, Bundibugyo ebolavirus: B8XCN0, Sudan ebolavirus: Q66814, Taï Forest ebolavirus: Q66810, Reston ebolavirus: Q66799. Sequence conservation was numbered according to EBOV GP and visualized using the Espript server (http://espript.ibcp.fr) and colored according to the percent equivalent scoring function with a cutoff of 70% (*38*).

#### Anti-GP mAb ELISAs

High-binding 96-well ELISA plates (Corning) were coated with 50 μL GP antigens in phosphate-buffered saline (PBS) at 4 μg/mL, and allowed to bind for 1 h at room temperature. After washing, the wells were blocked with PBS containing 3% bovine serum albumin (PBSA) for 1 h at room temperature, followed by washing then incubation with ADI-15946 or one of its mutant derivatives in serial dilutions of PBS. A horseradish-peroxidase conjugated anti-human secondary antibody (Santa Cruz Biotechnology) was added and allowed to bind for 1 h at room temperature and then subsequently detected by ultra-TMB (3,3′,5,5′-tetramethylbenzidine) substrate (ThermoFisher Scientific). Optical density was measured at 450 nm, and absorbance readings were subjected to a nonlinear regression analysis (GraphPad Prism software) to generate binding curves and calculate an EC_50_ value. ELISA assays were performed in triplicate, across seven 5-fold dilutions, beginning at 1000 ng/ml.

#### Biolayer Interferometry Assays

The Octet Red™ system (FortéBio, Pall) was used to determine the binding properties of different IgGs to various forms of EBOV GP. Anti-human Fc (AHC) capture sensors (ForteBio) were used for initial mAb loading at 25 mg/mL in 1X kinetics buffer (PBS supplemented with 0.002% Tween-20 and 1 mg/mL of BSA). Binding to GP was performed across two-fold serial dilutions of EBOV GPΔTM or GP_CL_. The baseline and dissociation steps were carried out in the 1X kinetics buffer as per the instrument manufacturer’s recommendations. Kinetic binding data in all cases are adequately and accurately described by a 1:1 binding model, but given the bivalent nature of the IgG (immobilized) and the trimeric state of GP (analyte), the association stoichiometry is likely to be more complex. Thus, k_off_/k_on_ likely reflects an ensemble of binding stoichiometries, and accordingly, we refer to this ratio as apparent K_D_ (K_d_^app^) throughout.

#### rVSV Neutralization Assays

Neutralization of rVSV: Recombinant vesicular stomatitis virus (VSV) expressing both enhanced green fluorescent protein (EGFP) and recombinant surface GP (rVSV-EBOV GP) in place of VSV G are previously described (*39*–*41*). Vero cells were seeded at 6.0 x 104 cells/well and cultured overnight in Eagle’s minimal essential medium (EMEM) supplemented with 10% fetal bovine serum (FBS) and 100 I.U./ml penicillin and 100 mg/ml streptomycin at 37°C and 5% CO_2_. The next day, virus was incubated with serial 3-fold antibody dilutions beginning at 350 nM (∼50 μg/ml) in serum free DMEM for one hour at room temperature before infecting Vero cell monolayers in 96-well plates. The virus was incubated with the cells in 50% v/v DMEM supplemented with 2% FBS, 100 I.U./ml penicillin and 100 μg/ml streptomycin at 37 °C and 5% CO_2_ for 14-16 hours before the cells were fixed and the nuclei stained with Hoescht. rVSV infectivity was measured by counting EGFP-positive cells in comparison to the total number of cells indicated by nuclear staining using a Cellinsight CX5 automated microscope and accompanying software (ThermoFisher Scientific).

#### Authentic Virus Neutralization Assays

Neutralization at BSL-4 was tested against replication-competent infectious EGFP-expressing EBOV and chimeric EBOV/BDBV-GP and EBOV/SUDV-GP constructs (referred as EBOV, BDBV and SUDV, respectively) in HTS format, as previously described (*42*). The neutralization assays were performed using Vero-E6 cells obtained from ATCC and maintained in Minimal Essential Medium (MEM) (ThermoFisher Scientific) supplemented by 10% fetal bovine serum (HyClone) and 1% penicillin-streptomycin at 5% CO2, 37°C. Neutralization assays were performed in triplicate, across twelve 4-fold dilutions, beginning from 200 μg/ml.

#### Authentic Virus Cooperativity Assays

The authentic Ebola virus/H.sapiens-tc/COD/1995/Kikwit-9510621 (EBOV/Kik-9510621; ‘EBOV-Zaire 1995’) (*43*), was used in this study. Antibodies were diluted to indicated concentrations in culture media and incubated with EBOV for 1 h. Vero cells were exposed to antibody/virus inoculum at an MOI of 0.2 plaque-forming unit (PFU)/cell for 1 h. Antibody/virus inoculum was then removed and fresh culture media was added. At 48 h post-infection, cells were fixed with formalin, and blocked with 1% bovine serum albumin. EBOV-infected cells and uninfected controls were incubated with EBOV GP-specific mAb KZ52 (*44*). Cells were washed with PBS prior to incubation with goat anti-human IgG conjugated to Alexa 488. Cells were counterstained with Hoechst stain (Invitrogen), washed with PBS and stored at 4°C. Infected cells were quantitated by fluorescence microscopy and automated image analysis. Images were acquired at 20 fields/well with a 20× objective lens on an Operetta high content device (Perkin Elmer, Waltham, MA). Operetta images were analyzed with a customized scheme built from image analysis functions available in Harmony software.

---

**Table S1.** Crystallographic data statistics.

|  | **EBOV GP<sub>CL</sub> – ADI-15946** |
| --- | --- |
| Resolution<sup>a</sup> (Å) | 48.54 – 4.10<br>(4.25 – 4.10) |
| Space group | P4<sub>3</sub>22 |
| Unit cell (Å)<br>(°) | 182.27 182.27 262.01<br>90 90 90 |
| Total Reflections<sup>a</sup><br>Unique Reflections<sup>a</sup><br>Reflections used in Refinement<sup>a</sup> | 463227 (46411)<br>35312 (3460)<br>35153 (3458) |
| Multiplicity<sup>a</sup> | 13.1 (13.4) |
| Completeness<sup>a</sup> (%) | 99.6 (100.0) |
| I/σ(I)<sup>a</sup> | 14.6 (1.8) |
| R<sub>merge</sub><sup>a,b</sup> | 0.20 (1.83) |
| R<sub>pim</sub><sup>a,c</sup> | 0.05 (0.46) |
| CC 1/2<sup>d</sup> | 1.00 (0.70) |
| Wilson B (Å<sup>2</sup>) | 159 |
| R<sub>work</sub><sup>e</sup> (%)<br>R<sub>free</sub><sup>f</sup> (%) | 26.2<br>28.1 |
| Reflections used in Refinement<sup>a</sup> | 35153 (3458) |
| RMSD<sup>g</sup> (bonds) (Å) | 0.01 |
| RMSD<sup>g</sup> (angles) (°) | 0.61 |
| Ramachandran favored (%)<br>Ramachandran outliers (%) | 98<br>0 |
| Average B-factor (Å<sup>2</sup>)<br>Macromolecules<br>Solvent | 114<br>114<br>107 |

<sup>a</sup> Values in parentheses are for the highest-resolution shell.
<sup>b</sup> $R_{\text{merge}} = \frac{\sum_{hkl} \sum_i |I_i(hkl) - \langle I(hkl) \rangle|}{\sum_{hkl} \sum_i I_i(hkl)}$
<sup>c</sup> $R_{\text{pim}} = \frac{\sum_{hkl} \{1/[N(hkl) - 1]\}^{1/2} \times \sum_i |I_i(hkl) - \langle I(hkl) \rangle|}{\sum_{hkl} \sum_i I_i(hkl)}$
<sup>d</sup> $CC = \frac{\sum (x - \langle x \rangle)(y - \langle y \rangle)}{[\sum (x - \langle x \rangle)^2 \sum (y - \langle y \rangle)^2]^{1/2}}$
<sup>e</sup> $R_{\text{work}} = (\sum_{hkl} \|F_{\text{obs}}\| - k \|F_{\text{calc}}\|) / (\sum_{hkl} \|F_{\text{obs}}\|)$ .
<sup>f</sup> $R_{\text{free}}$ is the same as $R_{\text{work}}$ with 5% of reflections chosen at random and omitted from refinement.
<sup>g</sup> RMSD, root mean square deviation.

**Table S2.**
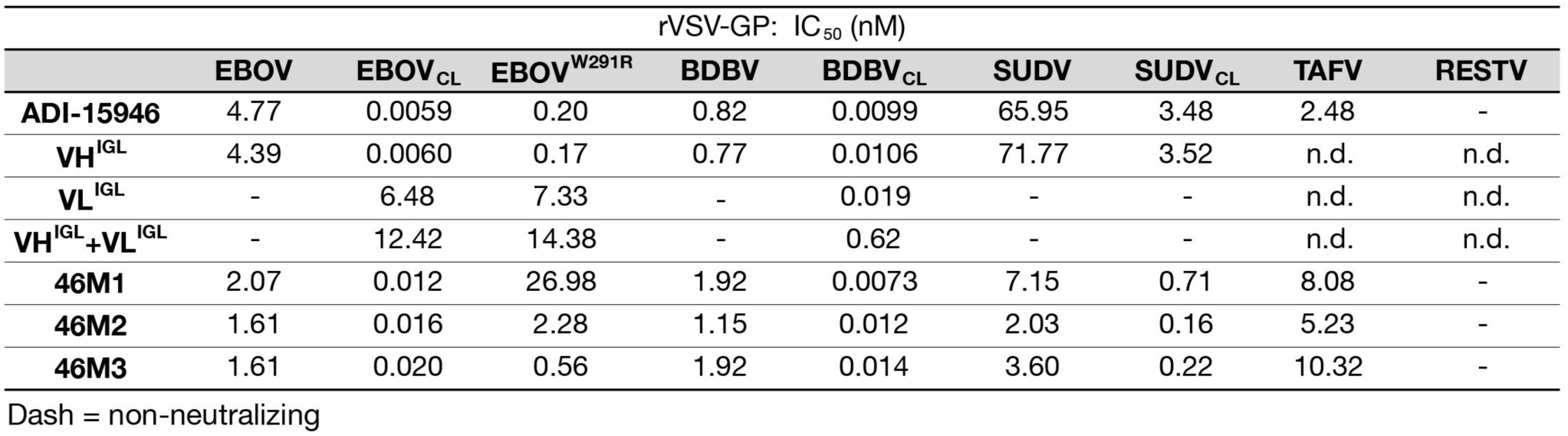
Neutralization potency (IC_50_) of ADI-15946 variants against rVSVs bearing filovirus glycoproteins.

**Table S3.**
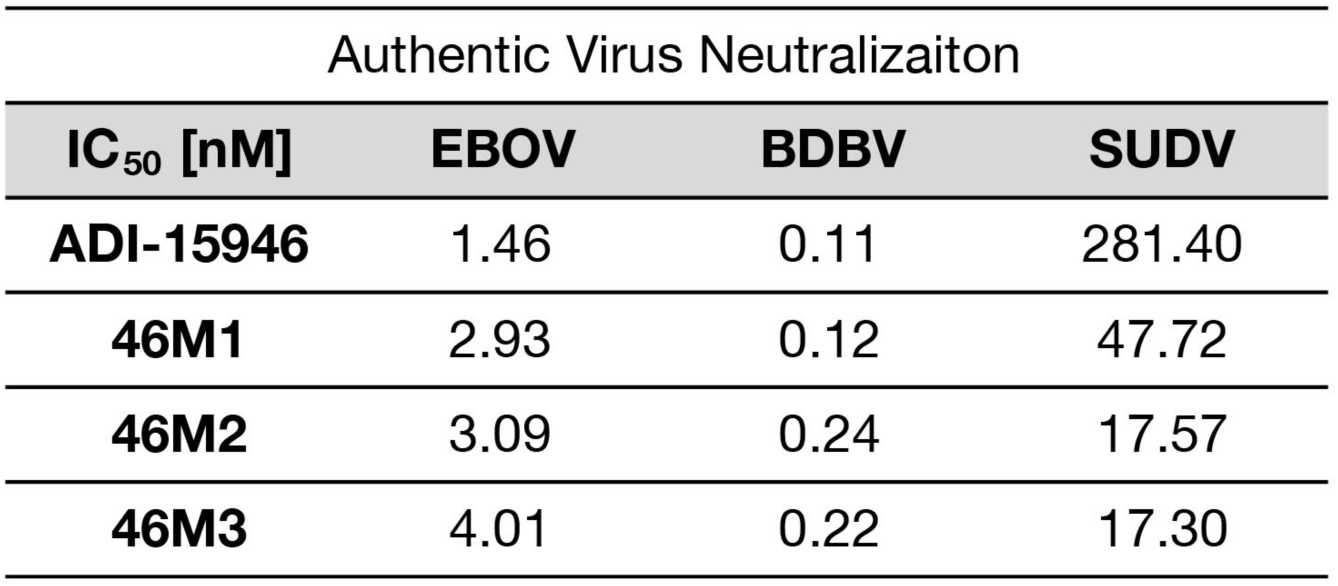
Neutralization potency (IC_50_) of ADI-15946 variants against authentic filoviruses.

| Authentic Virus Neutralization |  |  |  |
| --- | --- | --- | --- |
| IC<sub>50</sub> [nM] | EBOV | BDBV | SUDV |
| **ADI-15946** | 1.46 | 0.11 | 281.40 |
| **46M1** | 2.93 | 0.12 | 47.72 |
| **46M2** | 3.09 | 0.24 | 17.57 |
| **46M3** | 4.01 | 0.22 | 17.30 |

**Table S4.** Binding kinetics of ADI-15946 compared to inferred germline progenitors.

|  | KD (M) | KD Error | kon(1/Ms) | kon Error | kdis(1/s) | kdis Error |
| --- | --- | --- | --- | --- | --- | --- |
| Binding of ADI-15946 and its variants to EBOV GP full length at pH 7.5 |  |  |  |  |  |  |
| WT | 8.17E-09 | 6.27E-11 | 4.77E+04 | 2.79E+02 | 3.89E-04 | 1.93E-06 |
| HC<sup>IGL</sup> | 7.33E-09 | 1.02E-10 | 4.84E+04 | 4.90E+02 | 3.55E-04 | 3.41E-06 |
| LC<sup>IGL</sup> | 3.89E-08 | 3.55E-10 | 1.23E+04 | 9.47E+01 | 4.79E-04 | 2.36E-06 |
| HC<sup>IGL</sup>+LC<sup>IGL</sup> | 7.06E-08 | 4.20E-09 | 8.12E+03 | 4.79E+02 | 5.73E-04 | 4.25E-06 |
| Binding of ADI-15946 and its variants to EBOV GP cleaved at pH 5.5 |  |  |  |  |  |  |
| WT | <1.0E-12 | <1.0E-12 | 2.21E+06 | 1.22E+04 | <1.0E-07 |  |
| HC<sup>IGL</sup> | 4.59E-12 | <1.0E-12 | 2.20E+06 | 8.82E+03 | 1.01E-05 | 1.62E-06 |
| LC<sup>IGL</sup> | 3.59E-11 | <1.0E-12 | 2.24E+06 | 1.42E+04 | 8.04E-05 | 2.09E-06 |
| HC<sup>IGL</sup>+LC<sup>IGL</sup> | 1.07E-11 | 1.08E-12 | 2.14E+06 | 1.46E+04 | 2.29E-05 | 2.31E-06 |

**Figure S1.**
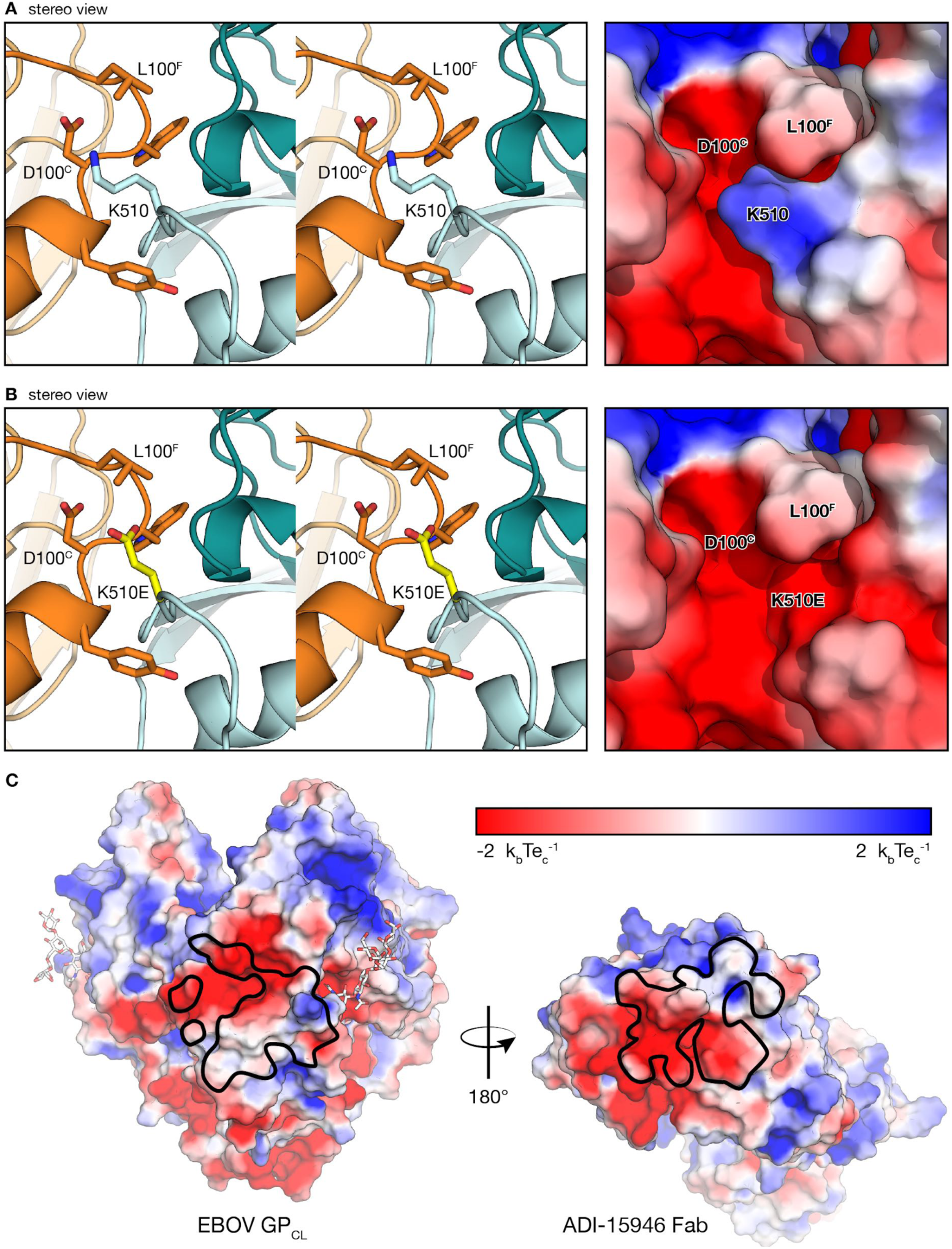
The K510E escape mutation likely clashes with ADI-15946 CDR H3. (A) Residue K510 of GP_2_ binds into a negatively charged pocket created by ADI-15946 CDR H3. Stereo views of the ADI-15946:GP_CL_ complex (ADI-15946 in orange, and GP_CL_ in dark and light teal for GP_1_ and GP_2_, respectively) are shown at left. The electrostatic surface potential is shown at right. (B) Modeling suggests that K510E, an escape mutant of ADI-15946, clashes with CDR H3 and introduces conflicting negative charge into the CDR H3 pocket. (C) Electrostatic surface potential generated using the APBS plugin with Pymol. The surface of GP_CL_ is shown on the left with the epitope outlined in black. The surface of ADI-15946 is shown on the right with the paratope outlined in black. All electrostatic surfaces are displayed with the same coloring as depicted in the included scale.

**Figure S2.**
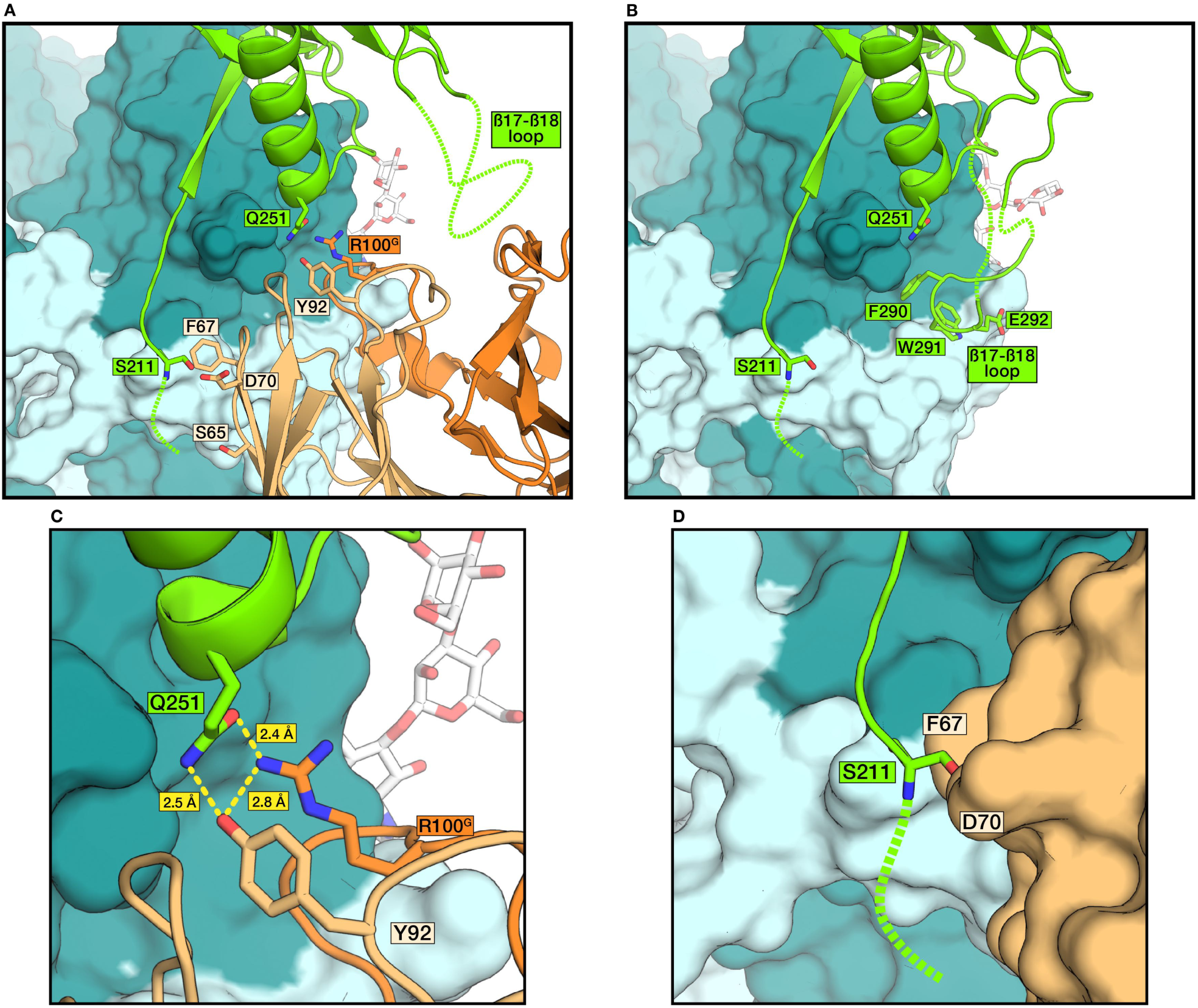
ADI-15946 likely inhibits cathepsin cleavage by blocking enzyme access to the β13-β14 loop. (A) Structural alignment of uncleaved GP to the ADI-15946:GP_CL_ complex shows that the mAb might make favorable interactions with the glycan cap even though binding displaces the glycan cap β17-β18 loop from the 3^10^ pocket. The surfaces of GP_1_ and GP_2_ are colored teal and light cyan respectively. ADI-15946 is shown in cartoon representation with the heavy and light chains colored dark orange and light orange respectively. The glycan cap (aligned from PDB 5JQ3) is shown as a light green cartoon. (B) The structure of uncleaved GP (PDB: 5JQ3, styled as in A) showing the position of the β17-β18 loop bound to the 3^10^ pocket. (C) The heavy and light chain of ADI-15946 may form favorable interactions with the glycan cap. For example, HC R100^G^ and LC Y92 are oriented such that they could potentially form hydrogen bonds with the 100% conserved GP_1_ residue, Q251. (D) Cathepsin access to the β13-β14 loop, shown as a green cartoon with the unresolved continuation of the loop as a dotted line, is likely inhibited by the steric bulk of ADI-15946 upon binding to GP. The structural alignment shows that the β13-β14 loop passes within close proximity to ADI-15946 and highlights a possible interaction between GP_1_ residue S211 and ADI-15946 LC residues F67 and D70.

**Figure S3.**
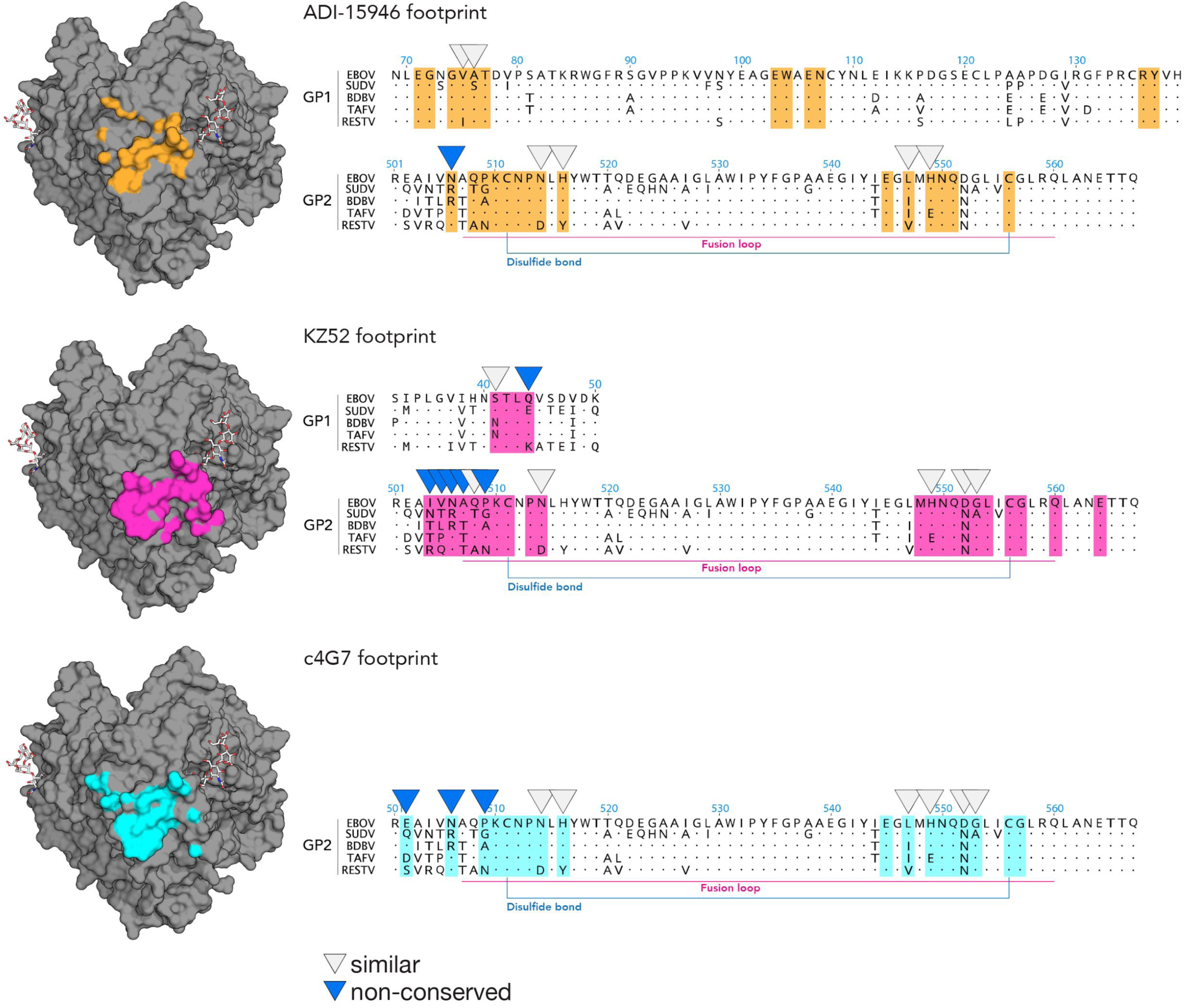
Comparison of the epitopes of ADI-15946 and monospecific base binders. In grey at left is a molecular surface of GP_CL_. Colored surfaces represent the respective footprints of ADI-15946 (orange), KZ52 (pink), and c4G7 (cyan). Next to each footprint is a partial amino acid sequence alignment of EBOV, SUDV, BDBV, TAFV and RESTV, with contact residues of each antibody highlighted in the appropriate color (ADI-15946 contacts are orange, KZ52 contacts are pink, etc.). Within each footprint, white inverted triangles indicate those contact residues that are similar across the ebolaviruses, while blue inverted triangles indicate those contact residues that differ more significantly across the ebolaviruses. Residues not indicated by triangles are either identical across the genus or not involved in the interface.

**Figure S4.**
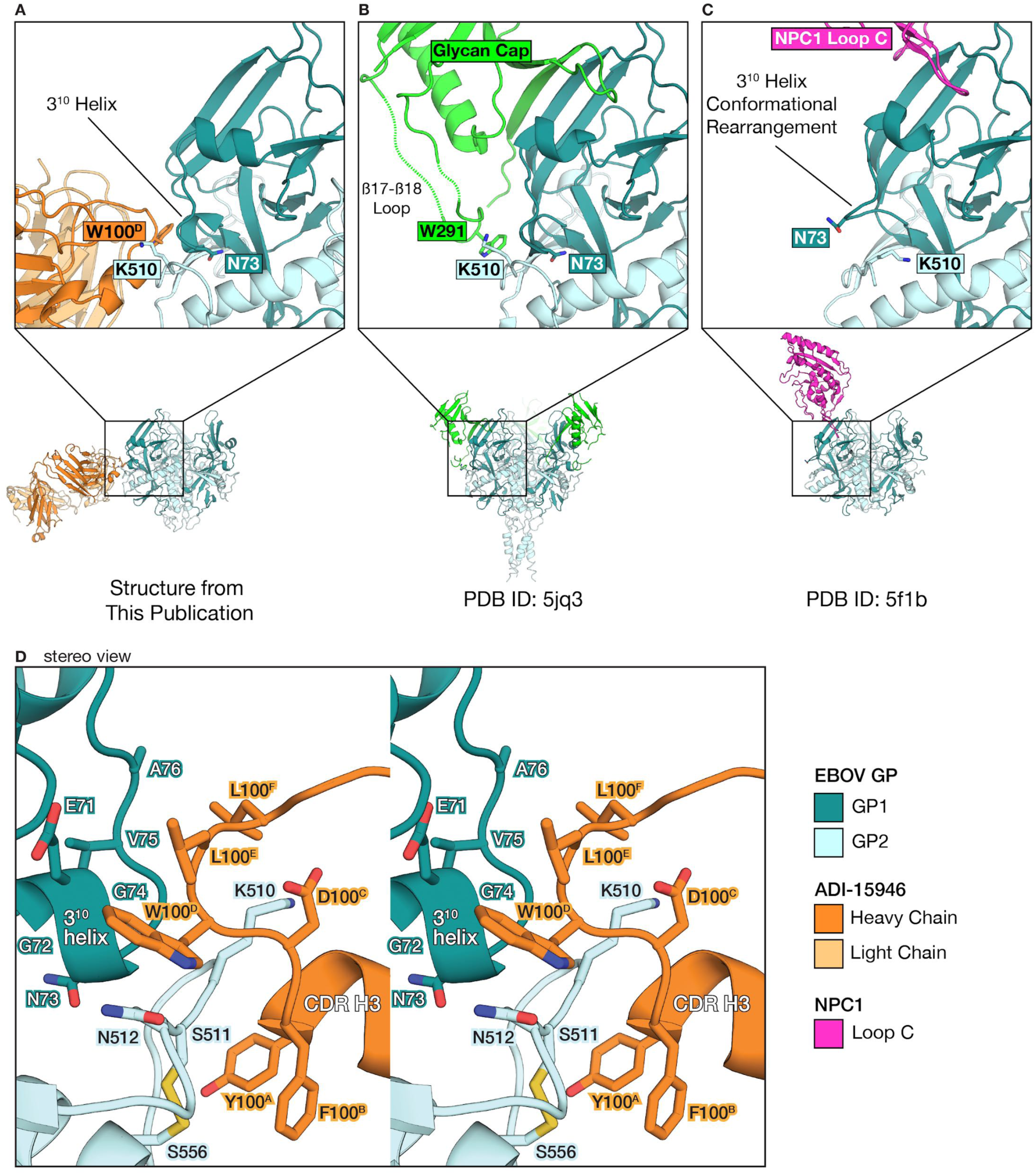
ADI-15946 CDR H3 binds to the 3^10^ pocket and may inhibit conformational rearrangements of the 3^10^ helix in the NPC1 bound complex. (A) ADI-15946 (orange) binds the 3^10^ pocket of GP that is usually occupied by the β17-β18 loop of the glycan cap (teal). (B) (C) Binding to NPC1 loop C (pink) induces a conformational change in GP: the 3^10^ helix unwinds and asparagine 73 (N73) becomes solvent exposed while lysine 510 (K510) inserts into the cavity left behind after the unwinding of the 3^10^ helix. ADI-15946 may prevent these conformational changes from occurring by locking down the 3^10^ helix with residues that mimic those of the β17-β18 loop. (D) Enlarged stereo view of ADI-15946 CDR H3 bound to the 3^10^ pocket showing interactions between CDR H3 sidechains and residues of the 3^10^ helix.

**Figure S5.**
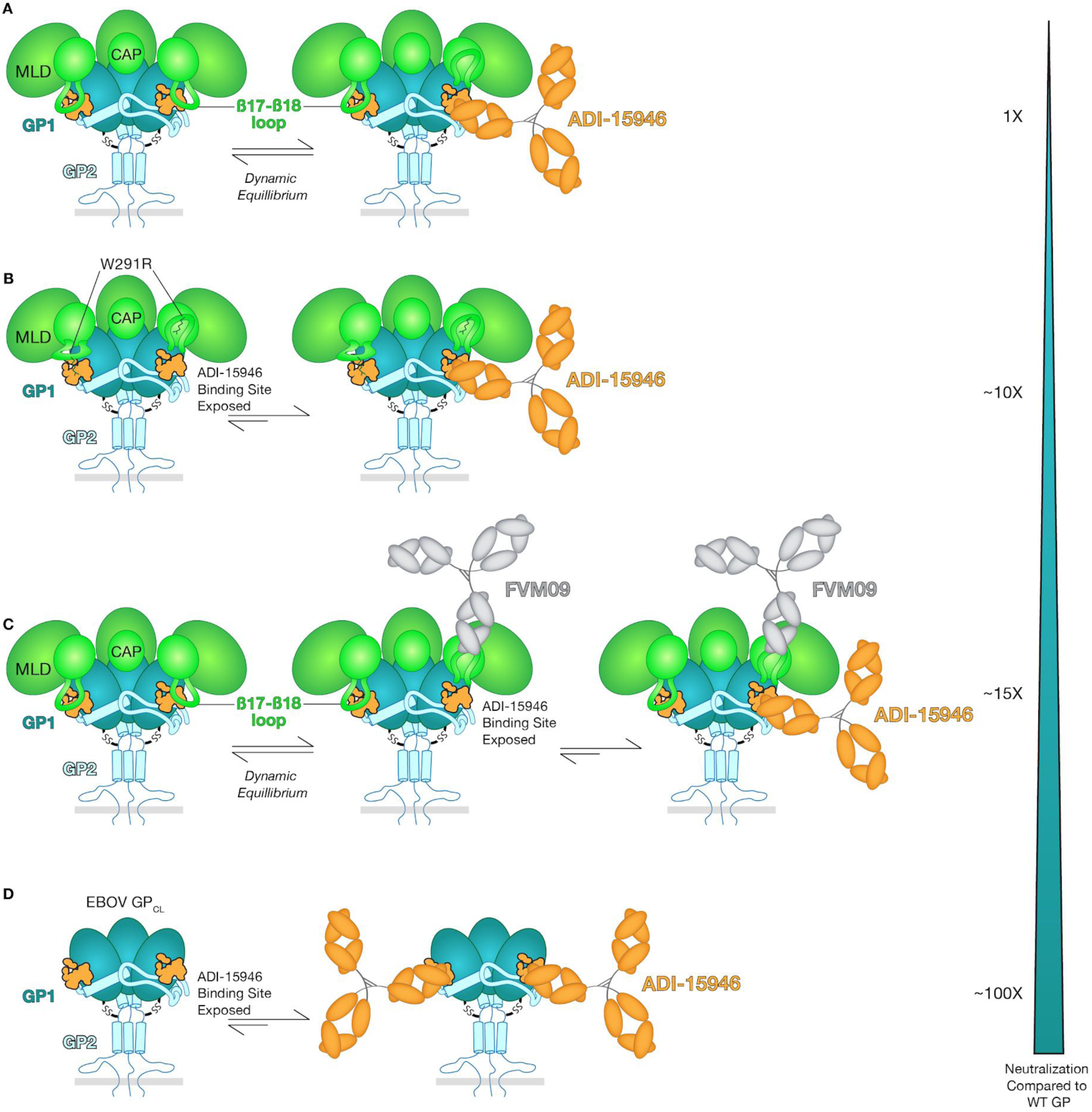
Conformational changes influence binding of ADI-15946. (A) Cartoon illustration of GP with mucin-like domains (MLD) and glycan cap domains (CAP) illustrated as green ovals. The β17-β18 loop of the glycan cap descends to cover the ADI-15946 epitope (orange). This loop may exist in a dynamic equilibrium between “tethered” and “loose” positions that mask and expose the ADI-15946 epitope. (B) The W291R point mutation in the β17-β18 loop results in enhanced exposure of the ADI-15946 epitope. (C) Antibody FVM09 (grey) which binds to the β17-β18 loop appears to lift it up and away (*23*), also better exposing the ADI-15946 epitope. (D) Enzymatic cleavage of GP deletes the glycan cap and the β17-β18 loop, better exposing the ADI-15946 epitope. Deletion of the glycan cap and β17-β18 loop enhances neutralization by ∼100-fold. Mutation of the loop and binding of the loop by FVM09 enhance ADI-15946 neutralization by ∼10- and 15-fold, respectively.

**Figure S6.**
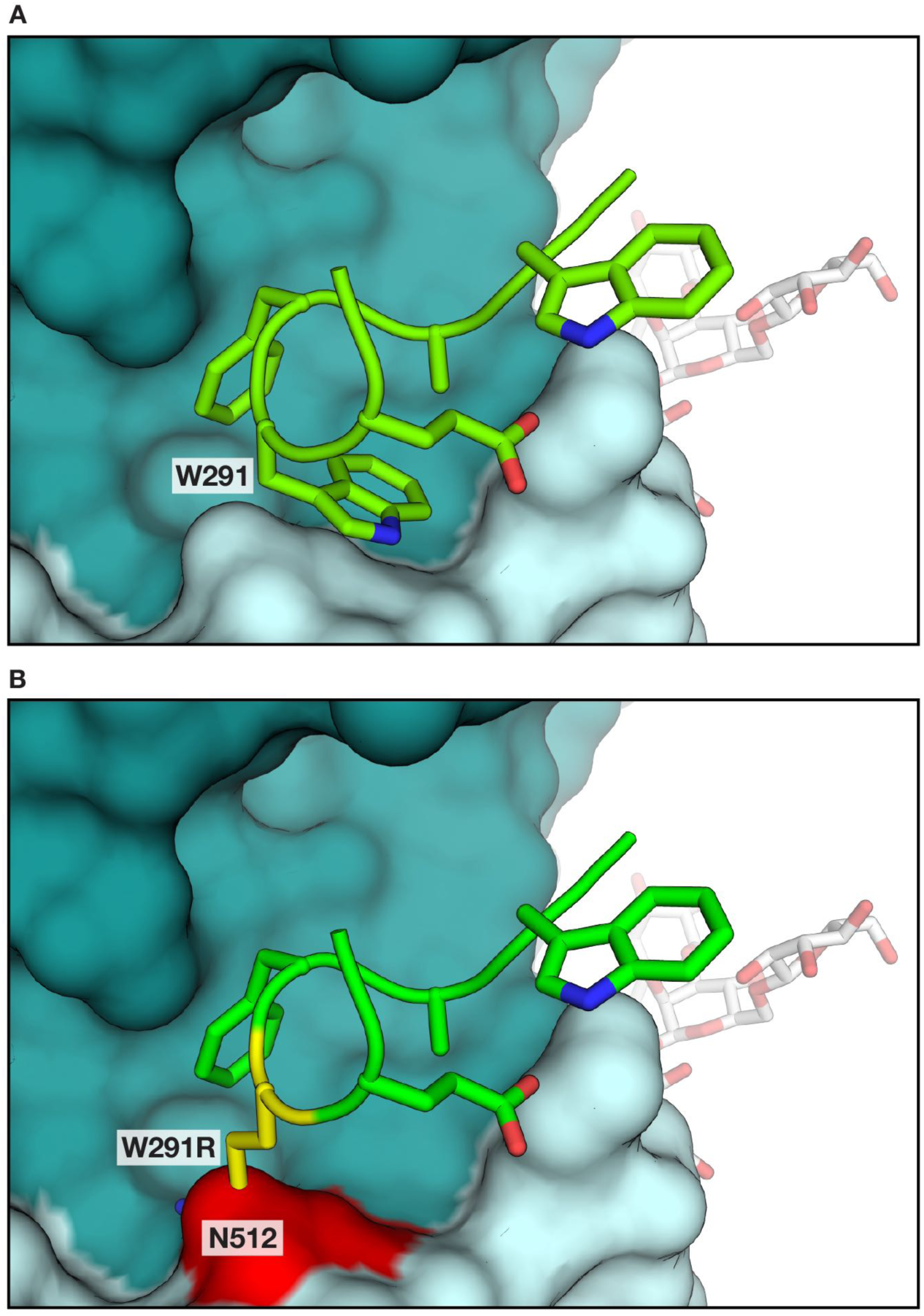
EBOV GP_1_ W291R mutation clashes with the 3^10^ pocket. (A) The position of the wild-type β17-β18 loop (green) on the GP core (teal molecular surface; PDB: 5JQ3). (B) The W291R mutation, modeled in yellow, introduces charge/steric clashes with EBOV GP_2_ N512 (shown as a red surface). The Arg residue was modeled using the most common rotamer, but many other possible Arg rotamers also clash with neighboring residues in the 3^10^ pocket. This mutation likely contributes to a decreased affinity of the β17-β18 loop for the 3^10^ pocket.

**Figure S7.**
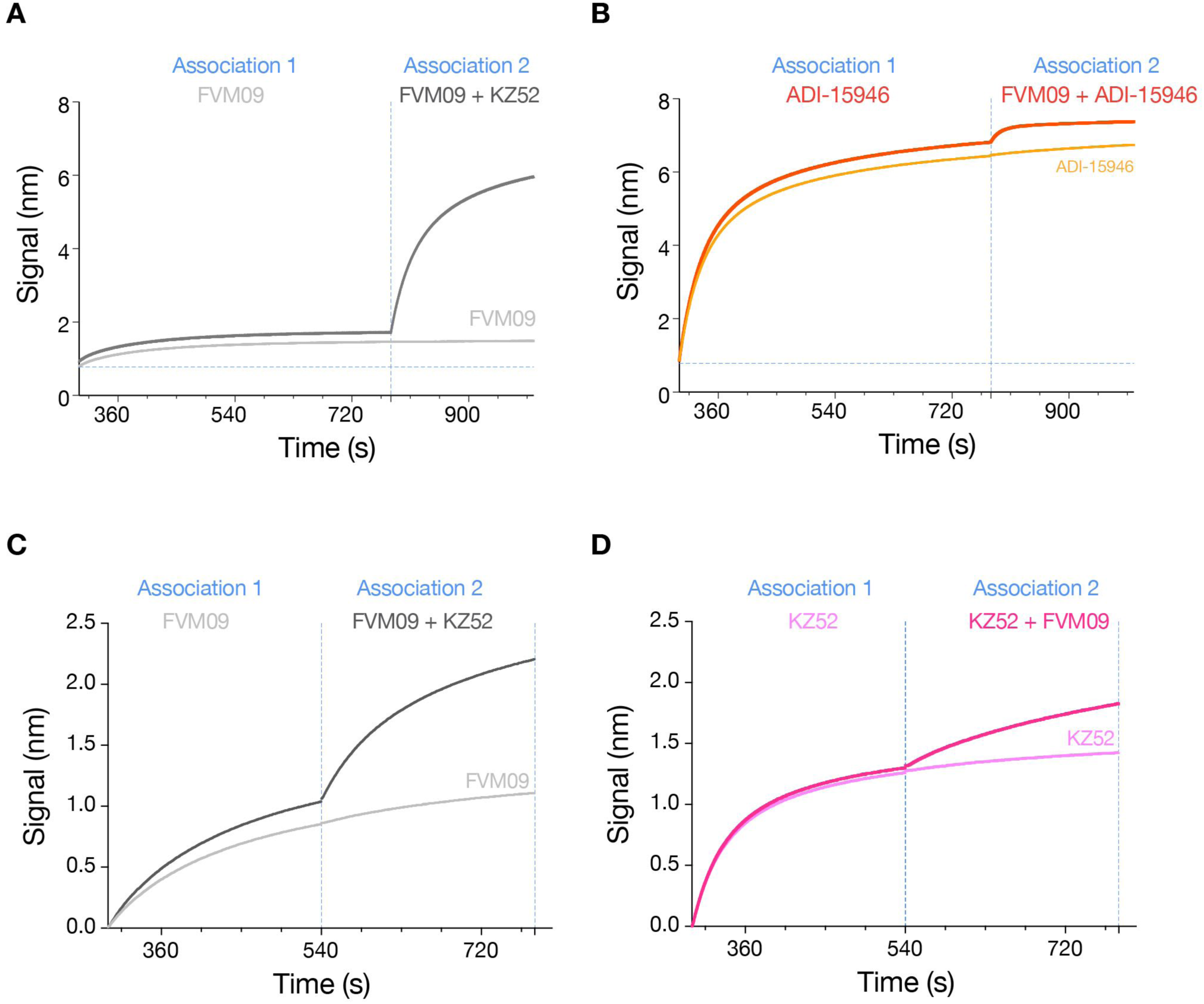
FVM09 competition biolayer interferometry assays. EBOV GP was loaded onto Nickel-NTA biosensors followed by two association steps of test antibodies. Pre-binding of FVM09 does not interfere with subsequent association of ADI-15946 (A) or KZ52 (C). In a converse experiment, pre-binding of ADI-15946 (B) or KZ52 (D) to GP does not interfere with subsequent binding of FVM09. In the second association steps, the first binder is maintained at the same concentration as in the first association step with addition of the competing antibody. All measurements were performed with antibodies at 330 nM concentration.

**Figure S8.**
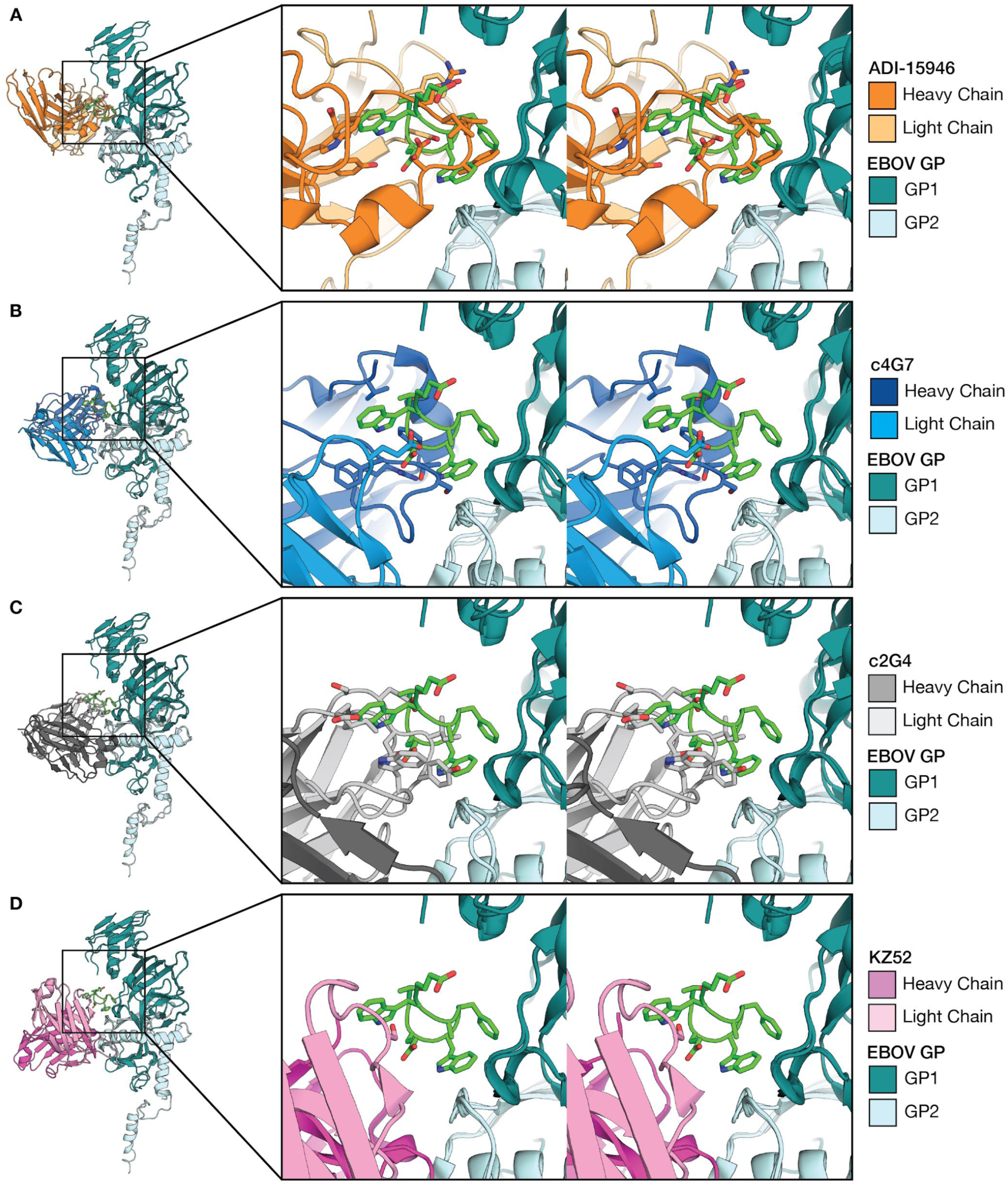
Clashes between the β17-β18 loop and ADI-15946, c2G4, c4G7, or KZ52. The cartoon representation on the left shows the various antibody’s Fragment variable (Fv) aligned to unbound, uncleaved EBOV GP (PDB: 5JQ3). The zoomed in panel shows a stereo view of each alignment. (A) The position of the β17-β18 loop (green) in unbound, uncleaved GP (PDB: 5JQ3) sterically overlaps the position of ADI-15946 in the ADI-15946:GP_CL_ crystal structure. (B–C) The position of the β17-β18 loop also interferes with the binding of c2G4 (PDB: 5KEL, blue), and c4G7 (PDB: 5KEN, grey). (D) The CDRs of KZ52 (PDB:3CSY, pink), however, do not clash with the β17-β18 loop, but rather form favorable interactions with the loop in its tethered position.

**Figure S9.**
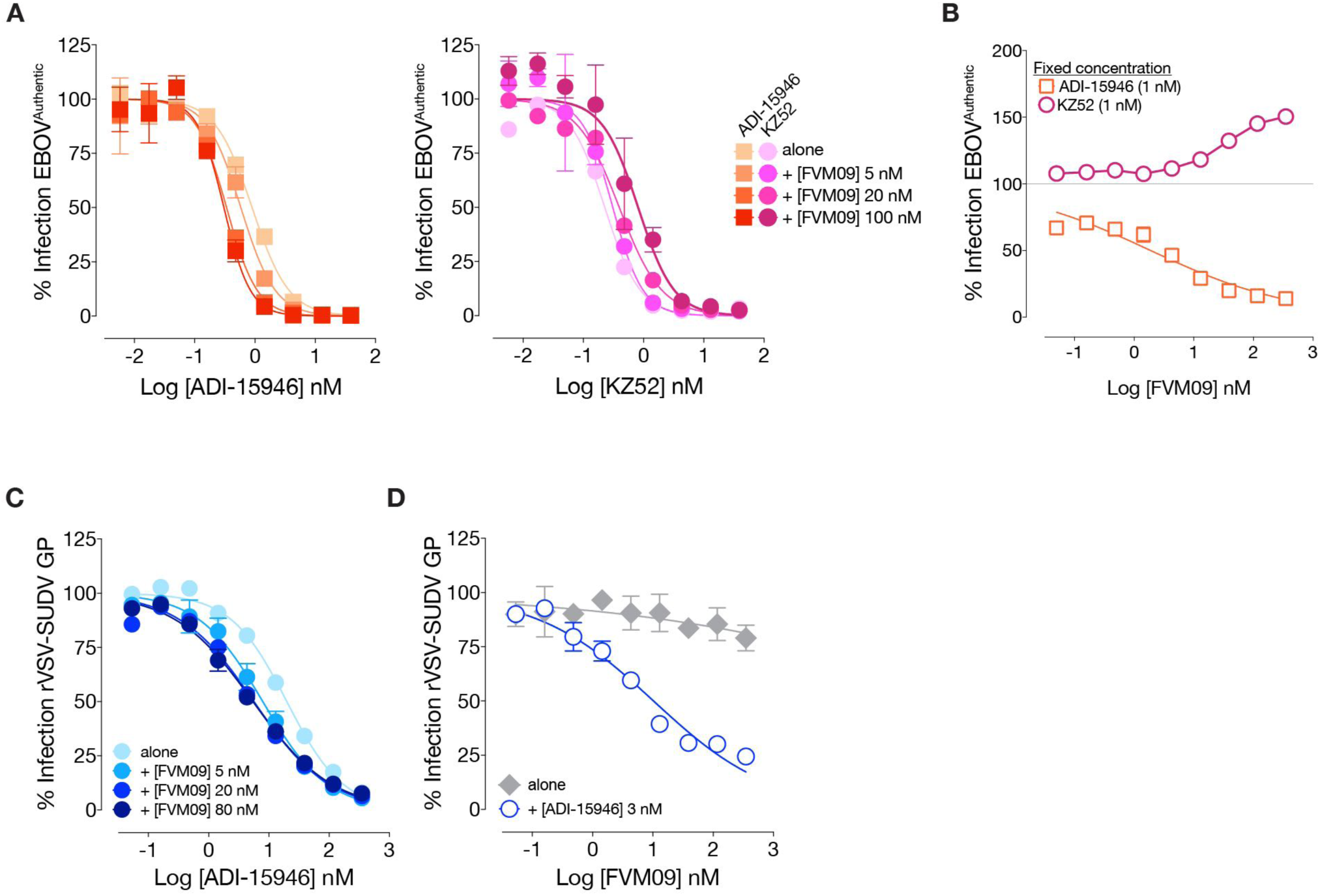
mAb FVM09 potentiates ADI-15946 neutralization of authentic EBOV. (A) Neutralization of Ebola virus by ADI-15946 (orange) and KZ52 (pink) was compared in the presence of increasing concentrations of the non-neutralizing FVM09 antibody (light to dark shading, 0-100 nM FVM09). Increasing amounts of FVM09 improved neutralization by ADI-15946, but slightly impaired neutralization by KZ52. Nine-point concentration series of ADI-15946 and KZ52 are plotted. (B) Neutralization of Ebola virus by a single 1 nM fixed concentration of ADI-15946 (orange squares) in the presence of increasing concentrations of FVM09. Neutralization by ADI-15946 is improved in the presence of increasing quantities of FVM09. KZ52, however, shows decreased neutralization in the presence of higher concentrations of FVM09. (C) Neutralization of SUDV GP-bearing rVSV by ADI-15946 in the presence of increasing concentrations of FVM09 (0-80 nM, light to dark blue). (D) Poor neutralization of rVSV-SUDV GP by FVM09 alone (grey filled diamonds), as compared to neutralization of rVSV-SUDV GP by 3nM ADI-15946 in the presence of FVM09 (blue open circles).

**Figure S10.**
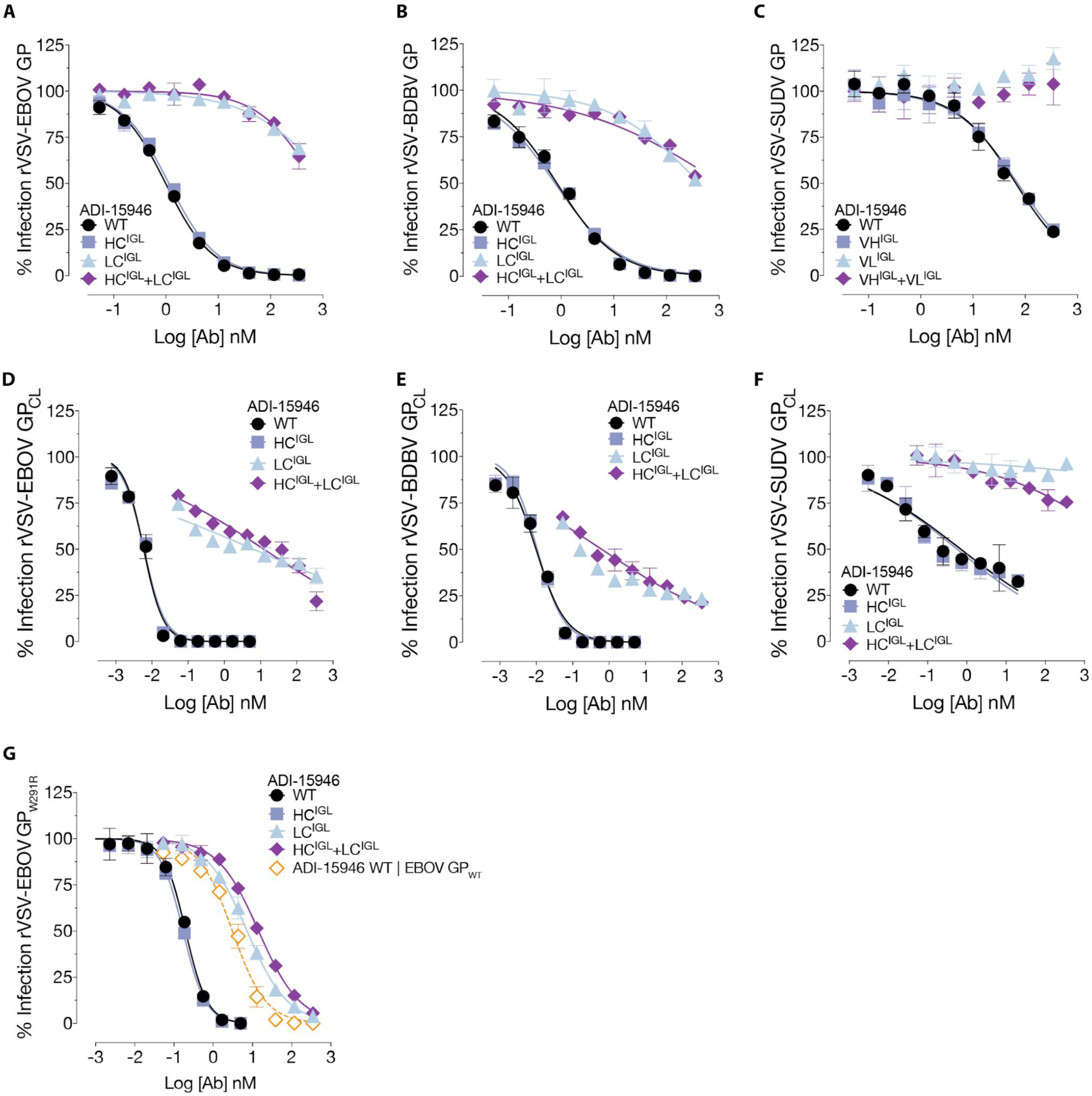
Germline neutralization data. (A-C) Neutralization of rVSV-EBOV GP, rVSV-BDBV GP, and rVSV-SUDV GP by wild-type ADI-15946 (black circles), heavy chain germline-revertant (HC^IGL^, grey squares), light chain germline-revertant (LC^IGL^; light blue triangles) and the HC^IGL^-LC^IGL^ combination (purple diamonds). (D-F) Neutralization of rVSV bearing GP_CL_ of EBOV, BDBV and SUDV, respectively. (G) Neutralization of rVSV-EBOV GP bearing a W291R point mutation by wild-type and germline revertant ADI-15946. For comparison, neutralization of rVSV-EBOV bearing uncleaved GP by wild-type ADI-15946 (orange open circles) is also shown. Virions bearing GP_CL_ are better neutralized by wild-type ADI-15946 and its heavy chain germline revertant (HC^IGL^; grey squares).

**Figure S11.**
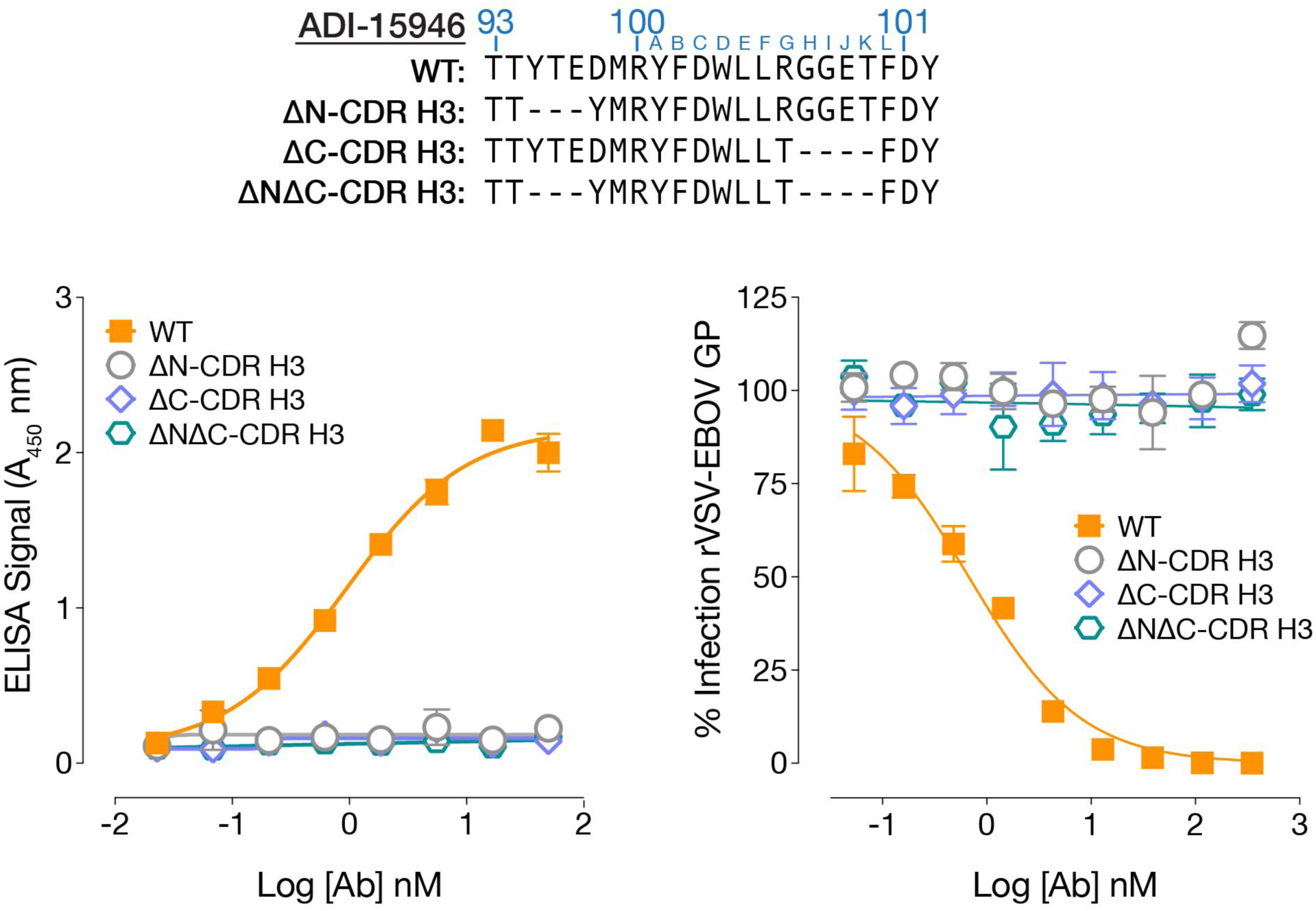
Truncations of CDR H3 abolish ADI-15946 activity. Truncated variants of ADI-15946 CDR H3 (removed residues shown as dashed lines in the alignment) were tested for binding (left panel) and neutralization (right panel) of rVSV-EBOV GP_FL_.

**Figure S12.**
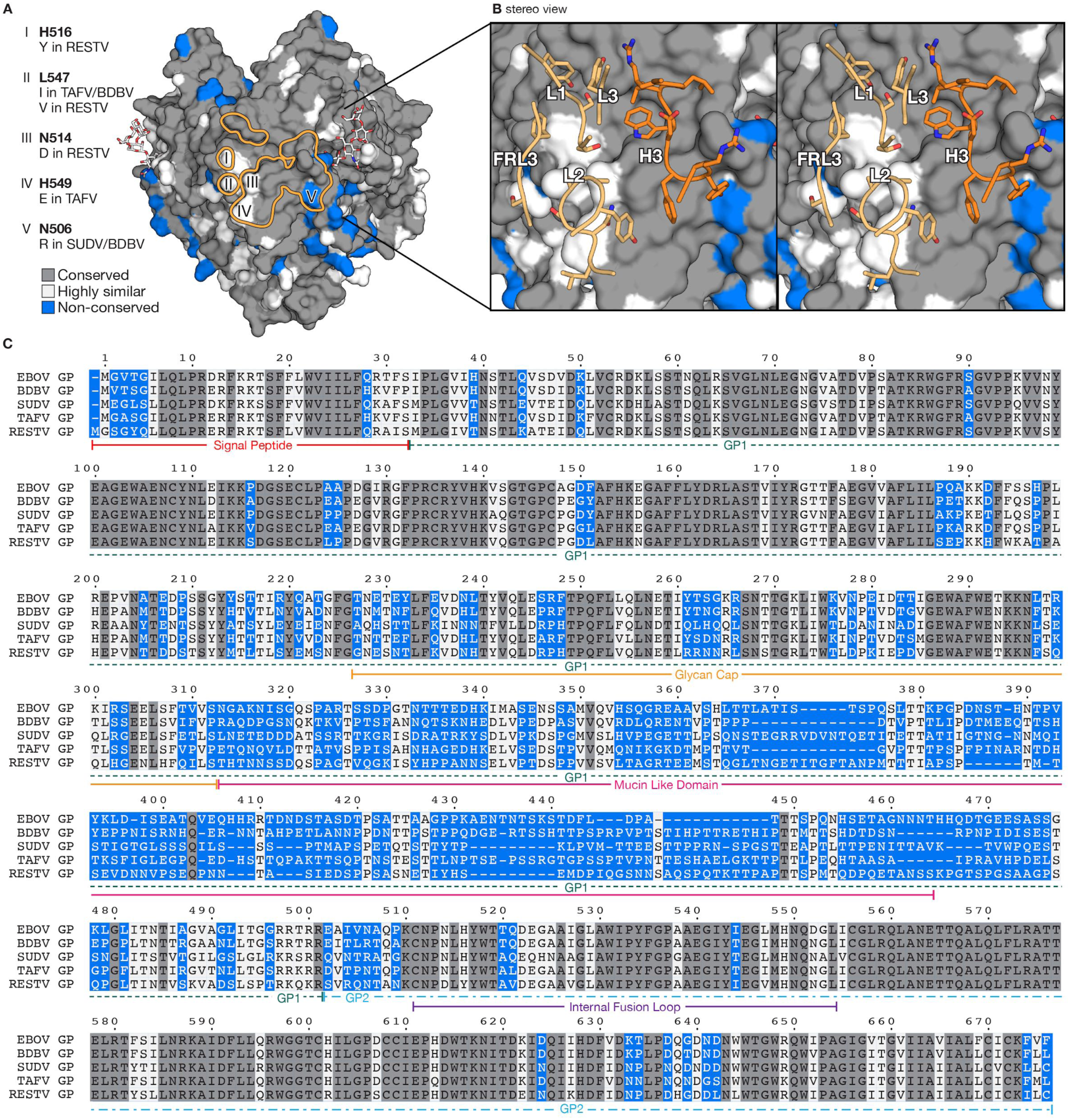
Ebolavirus conservation map and alignment. (A) Molecular surface of EBOV GP_CL_ with the ADI-15946 footprint outlined in orange. GP residues conserved across the ebolaviruses are in grey. Similar residues are in white and residues that differ significantly are in blue. Differences at five sites (I-V) are listed at left. (B) Stereo view of the ADI-15946:GP_CL_ interaction. GP is shown as a molecular surface and the interacting mAb residues are shown as sidechain residues along a ribbon backbone for clarity. The light chain is shown in light orange and the heavy chain is shown in darker orange. (C) Sequence alignment of GP from the five ebolaviruses, colored by conservation as in (A).

**Figure S13.**
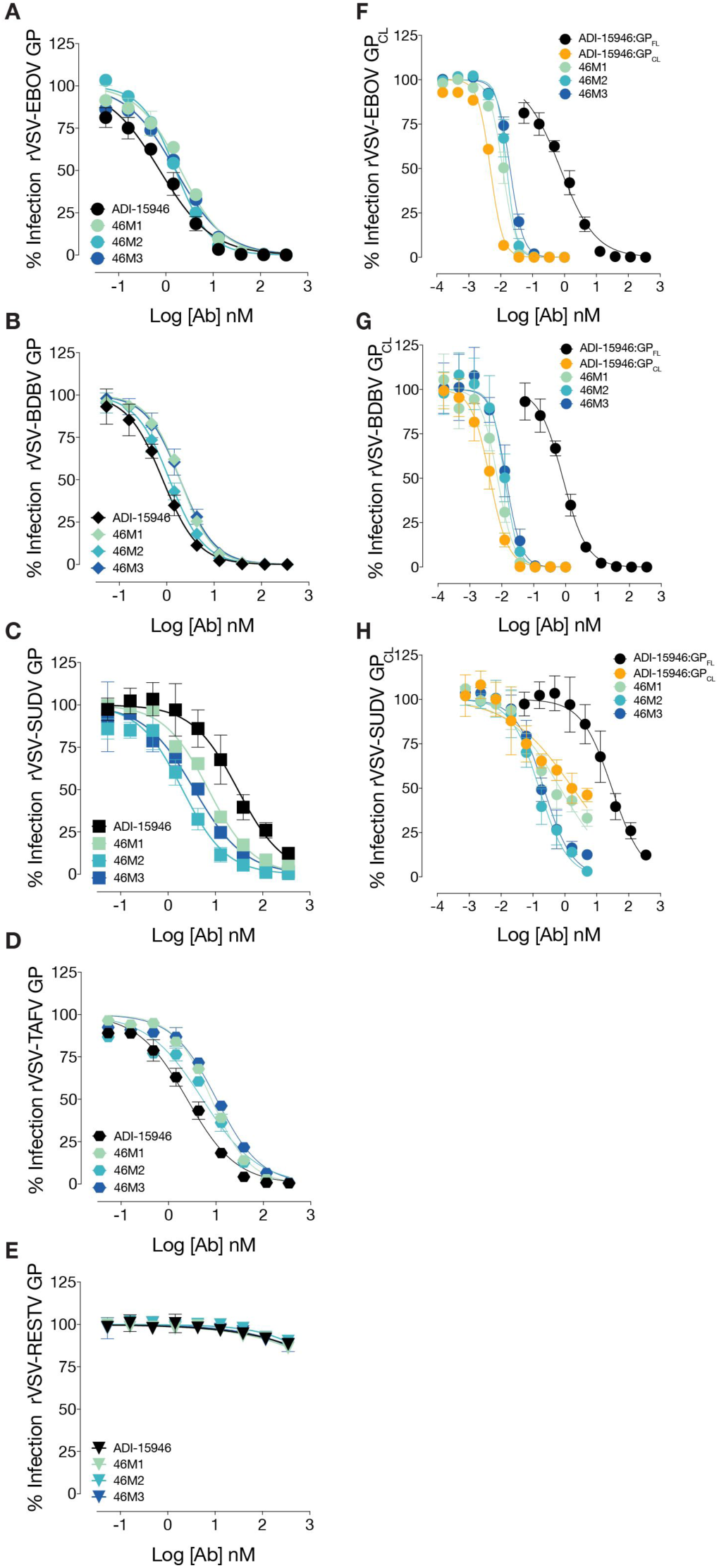
Neutralization of rVSVs bearing ebolavirus GPs by engineered variants of mAb ADI-15946. (A-E) Neutralization of rVSV bearing GP from EBOV, BDBV, SUDV, TAFV, and RESTV respectively. Wild-type ADI-15946 is in black, structure-guided mutants 46M1, 46M2 and 46M3 in light, medium and dark blue, respectively. (F-H) Neutralization of rVSV bearing GP_CL_ from EBOV, BDBV and SUDV, respectively. Neutralization of GP_CL_-bearing virions by wild-type ADI-15946 is in orange. Mutants 46M1-46M3 are in light to dark blue. Neutralization of virions bearing full-length GP by wild-type ADI-15946 is shown in black for comparison.

**Figure S14.**
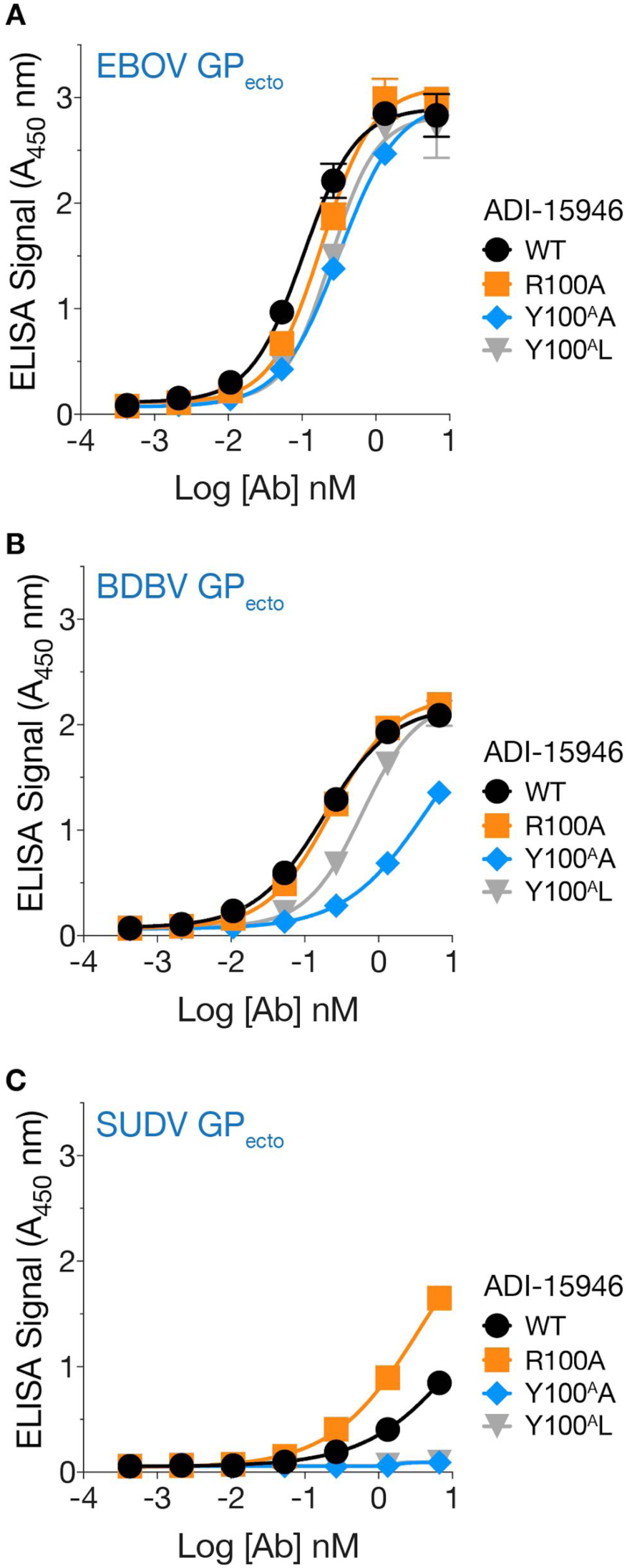
The R100A mutation in CDR H3 enhances ADI-15946 binding to SUDV GP_ecto_. Antibody variants were tested for binding to soluble EBOV (A), BDBV (B), or SUDV (C) GP ectodomain immobilized on ELISA plates.

**Figure S15.**
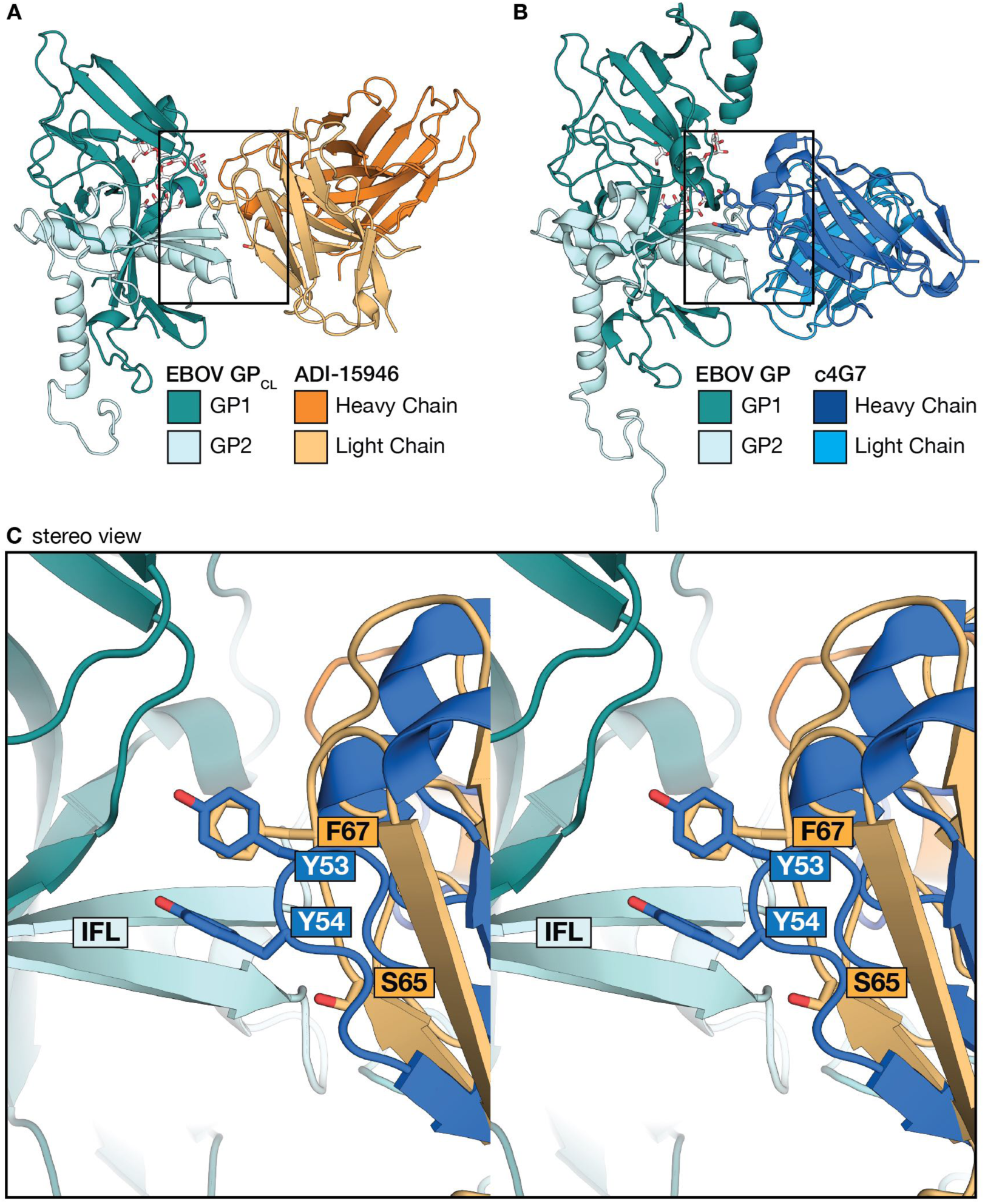
46M2 and 46M3 were designed to mimic interactions observed between c4G7 and EBOV GP. (A) ADI-15946 (orange, Fv shown) in complex with GP_CL_ (teal). (B) c4G7 (blue, Fv shown), in complex with uncleaved GP (PDB: 5KEN). (C) Zoomed in, superimposed stereo view of the ADI-15946 and c4G7 complexes (boxes shown in A and B). Both ADI-15946 and c4G7 position an aromatic residue (F67 for ADI-15946 light chain and Y53 for c4G7 heavy chain) along the IFL. The 46M2 and 46M3 variants of ADI-15946 were designed to mimic the binding of c4G7 with the residue substitutions S65Y and F67Y.

**Figure S16.**
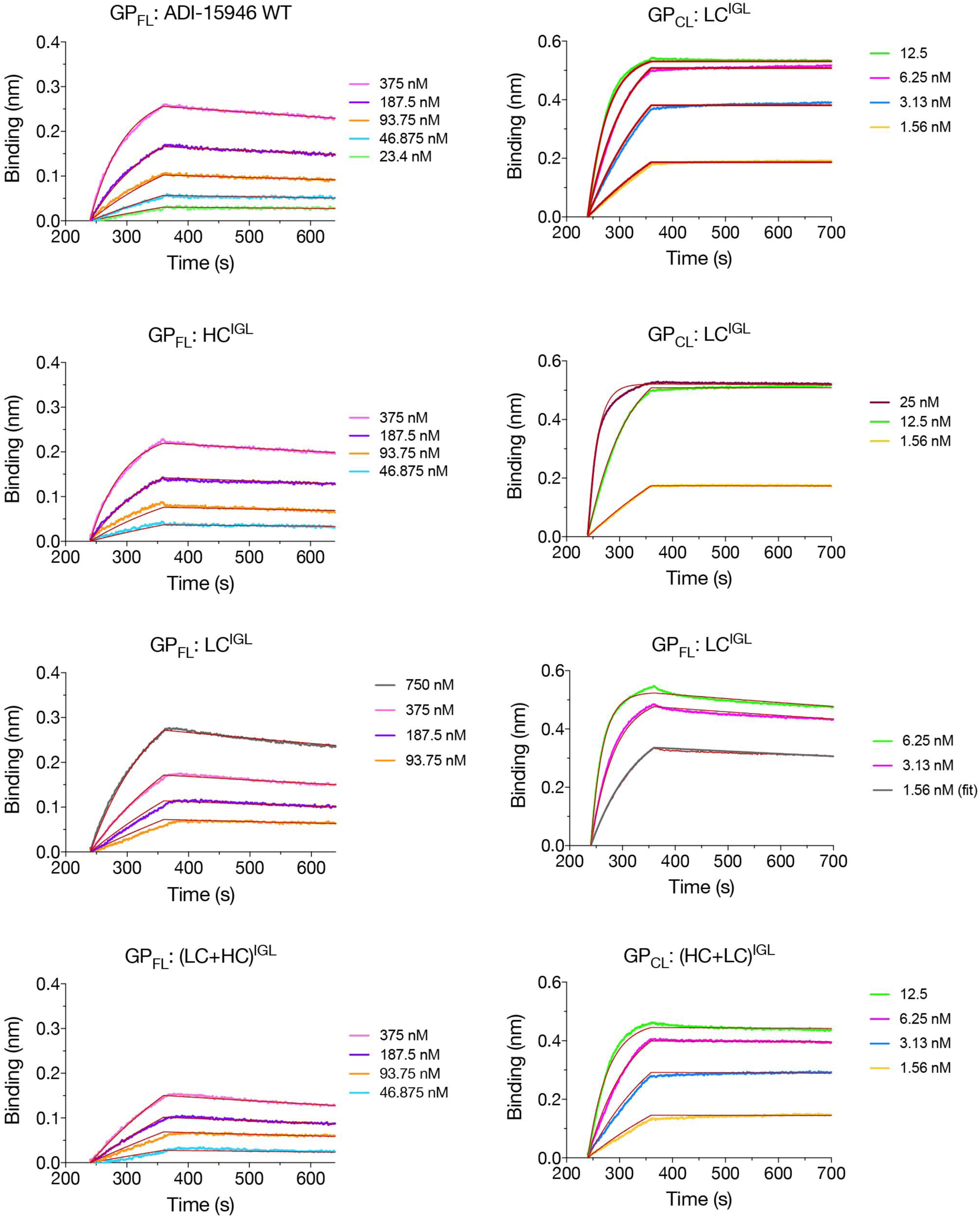
Binding kinetics showing that somatic hypermaturation of ADI-15946 LC improves binding to GP_FL_ and GP_CL_.

